# Hsp90-dependent triage of the glucocorticoid receptor via the CHIP E3 ubiquitin ligase

**DOI:** 10.1101/2025.05.28.656676

**Authors:** Cynthia M. Chio, Feng Wang, Chari M. Noddings, David A. Agard

## Abstract

The molecular chaperone Hsp90 is required to ensure the proper folding and activation of approximately 10% of the proteome throughout their cellular lifetime^1,2^. Given that Hsp90 recognizes and preferentially interacts with partially unfolded client states, it is well-poised to promote the degradation of misfolded clients and clients past the point of re-foldability^3,4,5^. Long-standing phenomenological data suggest that Hsp90 could directly present a client protein to an E3 ubiquitin ligase, in this way mediating degradative ubiquitylation of the client. However, this has yet to be directly demonstrated for any client. To assess this mechanism for the model Hsp90 client the glucocorticoid receptor (GR), a ligand-dependent transcription factor, we reconstituted *in vitro* the Hsp90-dependent folding cycle for GR^6^, in the presence of the E3 ligase CHIP. Remarkably, we discovered that CHIP binds to and efficiently ubiquitylates GR as presented by a previously characterized Hsp90-dependent pro-folding state called the GR-maturation state (Hsp90-p23-GR), in which GR has been visualized via cryo-EM to be folded in the native state and ligand-bound^7^. We determined a low-resolution (∼14 Å) cryo-EM reconstruction of the four-component Hsp90-p23-GR-CHIP complex, and furthermore resolved from the same dataset a 3.53 Å reconstruction of a Hsp90-p23-GR complex in which, surprisingly, GR is partially disordered and ligand-free—in contrast with what we previously observed^7^, presumably as a consequence of interaction with CHIP. Our results suggest a mechanism by which Hsp90 can simultaneously partition a client between both folding and degradation outcomes and in this way directly mediate protein-triage decisions through its interactions with E3 ligases.

## Introduction

A fine balance between folding and degradation is critical for protein homeostasis and needs to be precisely regulated. Molecular chaperones have long been recognized as essential players supporting protein folding in the cell^8^. Their function in simultaneously mediating protein degradation—in so doing, triaging proteins between life and death—has also been suggested^9,10,4,5^, and indeed, recently, this was demonstrated directly for the first time in a fully reconstituted chaperone system for the pathway of tail-anchored membrane protein insertion^11^.

Among other chaperones at large, the ATP-driven Hsp90 is unique in acting in the late stages of folding on near-native states of its many ‘client’ proteins^1^. Focused efforts over the past decade have delivered atomic-resolution mechanistic insights into how Hsp90 promotes client folding^6,12,13,7,14,15,16,17^. Lagging much farther behind is an understanding of how Hsp90 might facilitate client degradation. It has long been known that chemical inhibition of the Hsp90 ATPase leads to rapid client degradation in cells^18,19,20,21,22,23,24,25^. While this had initially been thought of as a consequence of removing Hsp90, it later became apparent that these inhibitors can stall the chaperone ATPase cycle in what has been called a “client loading” complex—in which the client could remain significantly unfolded while stabilized by Hsp90—and prevent progression to a ”client maturation” complex—in which the client is better folded by Hsp90^6,13,7^. In the context of the older phenomenological data above, these more recent biochemical observations recommend a hypothesis where, upon intercepting a client protein, Hsp90 might naturally promote either folding or degradation outcomes, depending partly on the lifetime of Hsp90-client association, which Hsp90-inhibition would artificially prolong. In this way, Hsp90 naturally actively mediates protein-triage decisions, in support of protein quality control. Yet the mechanism by which Hsp90 might connect client to a degradation outcome remains poorly defined. There is, however, a wealth of data linking Hsp90 to ubiquitin-dependent degradation pathways. Specifically, Hsp90 has been found to interact with or co-immunoprecipitate with a wide variety of E3 ubiquitin ligases^26,27,28,29,30,31,32^, whose ubiquitylation activity on protein substrates can mark a substrate for degradation by the ubiquitin-proteasome system^33,34^. Together, these lines of evidence support a mechanistic model in which Hsp90 facilitates proteasomal client degradation by directly presenting a stalled client protein to an E3 ligase, forming an Hsp90-client-E3 ligase “client triage” complex that promotes client ubiquitylation. However, direct demonstrations of such an Hsp90- dependent triage mechanism have been lacking to date.

In the simplest model of such Hsp90-dependent triage, Hsp90 states on-pathway to client folding are in fact client-triage states capable also of recruiting E3 ligases and allowing for client ubiquitylation, with kinetic control determining the net balance between folding and ubiquitylation/degradation outcomes. To test this hypothesis, we have turned to the glucocorticoid receptor (GR) client system, a ligand-dependent transcription factor and constitutive Hsp90 client. We previously showed it was possible to fully reconstitute the Hsp90-mediated GR folding cycle *in vitro* using a set of individually purified factors beyond Hsp90 alone^6^, permitting controlled, modular access to multiple distinct chaperone-GR states that could potentially serve as client- triage points (**Fig. 1a**). The folding cycle begins with another molecular chaperone Hsp70 and its cochaperone Hsp40 partially unfolding GR, specifically inactivating GR ligand-binding. Next, in the presence of the cochaperone Hop, Hsp70-GR is loaded onto Hsp90, forming a heterochaperone Hsp90-GR-Hsp70-Hop “client loading” state. In this state, GR remains significantly unfolded, cradled by Hsp90, Hsp70, and Hop^13^. Finally, partial ATP hydrolysis by Hsp90 triggers the release of Hsp70 and Hop, and following incorporation of the cochaperone p23, a Hsp90-p23-GR “client maturation” state is formed. In the maturation state, with Hsp70 and Hop gone, GR is refolded and reactivated for ligand binding^7^. In short, whereas Hsp70 unfolds GR, Hsp90—depending on its ATP-hydrolysis state—has the capacity to either stabilize the unfolded GR or re-fold the client; the generality of this Hsp70-Hsp90 antagonism has been shown for other client systems^35,36,37^. We hypothesized that these Hsp90-folding states could in fact serve as client-triage points in the presence of a triage E3 ligase and relevant ubiquitylation machinery, accommodating E3 incorporation and promoting efficient GR ubiquitylation upon binding by the E3 enzyme.

**Figure 1:**
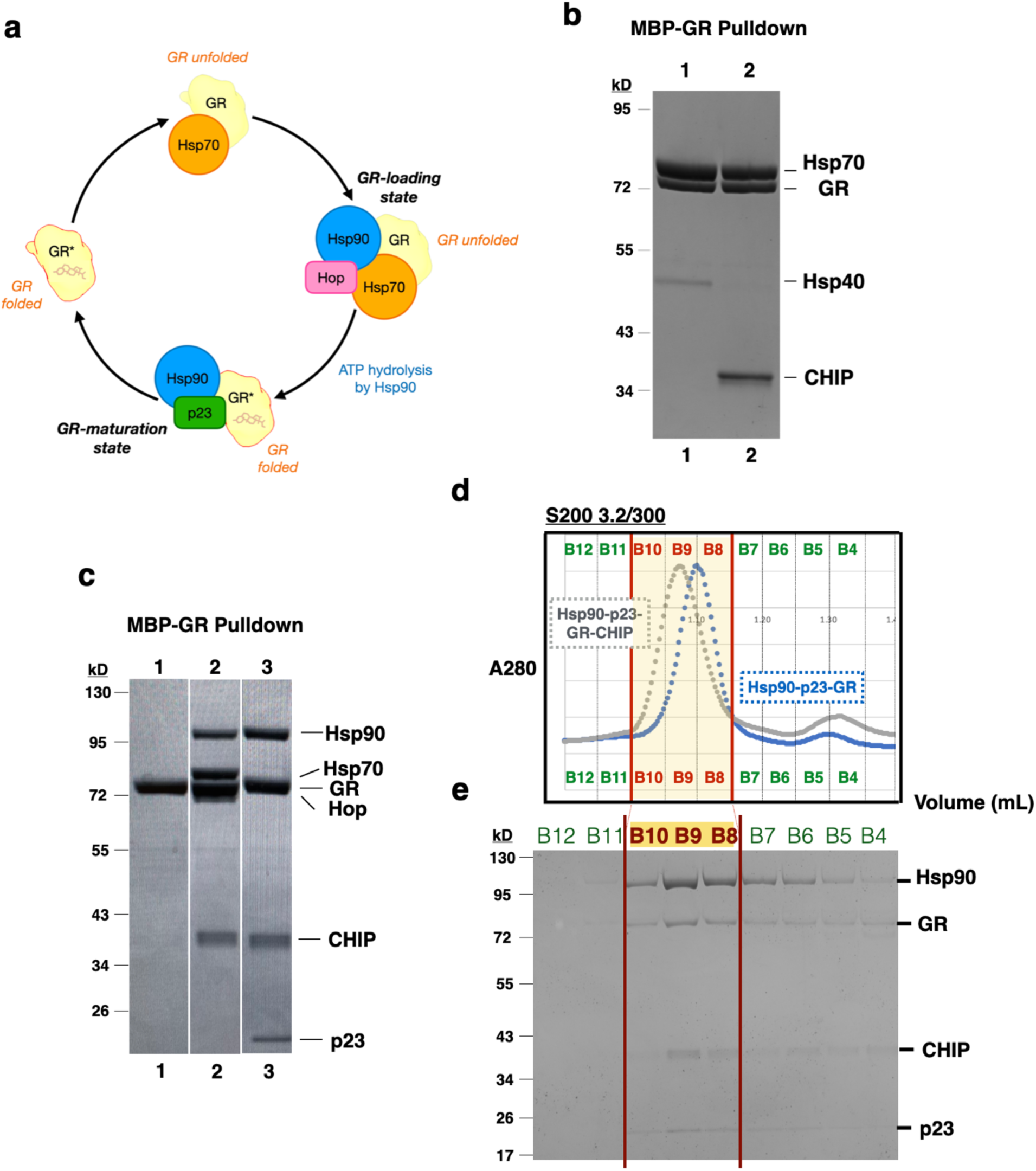
CHIP incorporates into the GR-loading complex (Hsp90-GR-Hsp70-Hop) and the GR-maturation complex (Hsp90-p23-GR) **a**, Ligand binding by GR is regulated by the ATP-driven molecular chaperones Hsp70 and Hsp90. GR loses its ligand-binding ability upon association with Hsp70 but regains it upon association with Hsp90 and its cochaperone protein p23 in a “GR-maturation state” (Hsp90-p23- GR). Formation of this latter complex is preceded by an obligate intermediate called the “GR- loading state” (Hsp90-GR-Hsp70-GR) in which inactive GR is in complex with both Hsp70 and Hsp90 as well as a cochaperone protein Hop. Shown is a representation of just the ligand- binding domain of GR (GR LBD). **b**, Coomassie-stained SDS-PAGE with elution from MBP-GR pulldown from the *in vitro* reconstituted GR chaperone cycle. Assay conditions are as follows - Lane 1: 5uM MBP-GR + 2uM Hsp40, 15uM Hsp70, 5mM ATP. Lane 2: 5uM MBP-GR + 2uM Hsp40, 15uM Hsp70, 15uM CHIP, 5mM ATP. **c**, Coomassie-stained SDS-PAGE with elution from MBP-GR pulldown from the *in vitro* reconstituted GR chaperone cycle. Assay conditions are as follows - Lane 1: 5uM MBP-GR + 15uM CHIP; Lane 2: 5uM MBP-GR, 2uM Hsp40, 15uM Hsp70, 15uM Hop, 15uM D93N Hsp90, 20uM CHIP, 5mM ATP; Lane 3: 5uM MBP-GR, 2uM Hsp40, 5uM Hsp70, 5uM Hop, 15uM Hsp90, 15uM Bag-1, 15uM p23, 20uM CHIP, 5mM ATP, 20mM molybdate. **d**, S200 3.2/300 gel-filtration profiles of the GR maturation complex (blue) and Hsp90-p23-GR-CHIP (grey) purified by MBP-GR pulldown from the reconstituted GR chaperone cycle. **e**, Coomassie-stained SDS-PAGE of elution fractions from gel filtration of Hsp90-p23-GR-CHIP.

While many E3 ligases complex with Hsp90 (as cited previously), the CHIP E3 ligase^38,29^ is a particularly attractive candidate for a triage E3 as it contains a tetratrico<u>p</u>eptide repeat (TPR) domain, a conserved domain that directly engages the EEVD peptide motif present at the C-termini of Hsp90 and Hsp70. Overexpression of CHIP has also been observed to shorten GR’s half-life in the cell^40^, suggesting GR to be a substrate of CHIP. Given its ability to engage both Hsp90 and Hsp70, CHIP would seem well-poised to intercept chaperone-GR complexes sampled along the folding cycle and facilitate GR ubiquitylation. Indeed, CHIP-mediated ubiquitylation of GR has been demonstrated on the receptor population that co-immunoprecipitates with Hsc/Hsp70^41^, implicating the existence of an Hsp70-GR-CHIP triage complex. Notably, the presence of Hsp90 was not examined in these semi-purified complexes. As such, the ability of the GR-loading state and the GR-maturation state to serve as client-triage points in the presence of CHIP remains unexplored.

Indeed, as a potential partner for chaperone-mediated protein degradation, historically CHIP has most often been considered in the context of its interaction with Hsp/Hsc70 rather than Hsp90^42^. As with GR, CHIP-mediated ubiquitylation of the Hsp90 clients p53 and the cystic fibrosis transmembrane conductance regulator (CFTR) has been investigated largely or only, respectively, in the presence of Hsc70 machinery^43,44,45^. In examples where Hsp90-dependent CHIP activity has been probed, overly reductionist *in vitro* mixes combining only Hsp90, client, and CHIP have been explored^46,42,47^, which are likely to fall short of reconstituting bona fide Hsp90-client complexes given the field’s emerging understanding of the full chaperone/co-chaperone requirements for faithfully recapitulating Hsp90-client complexes—as described above for GR and corroborated elsewhere for other client systems^6,35,36,37^. Indeed, this could account for results showing Hsp90-dependent client ubiquitylation to be less robust than Hsp70-dependent client ubiquitylation^42^. Given our now deeper understanding of the GR-chaperone cycle, there is the possibility that GR could also be efficiently presented to CHIP via either the client loading complex (Hsp90-GR-Hsp70-Hop) or the client maturation state (Hsp90-p23-GR).

Here we report, to our knowledge for the first time, that CHIP can intercept the GR-maturation state and strikingly subvert it for a degradation outcome, efficiently and preferentially ubiquitylating GR compared to GR presented by Hsp70 alone. We report also a low-resolution (∼14 Å) single-particle cryogenic electron microscopy (cryo-EM) reconstruction of CHIP bound to the maturation state. In a subset of particles within this dataset yielding a 3.53 Å reconstruction of Hsp90-p23-GR, GR surprisingly appears ligand-free and partially disordered. This is in stark contrast with the well-folded, ligand-bound GR structure our lab previously resolved in the maturation state complex^7^. This suggests CHIP preferentially recognizes partially unfolded chaperone-bound substrates in the context of Hsp90 and efficiently ubiquitylates them. Our work calls for an update of Hsp90 function to include client presentation for degradation.

## Results

### CHIP integrates into the GR-loading and GR-maturation complexes

To test whether CHIP can intercept chaperone-GR complexes sampled along GR’s folding cycle (**Fig. 1a**), we staged the three distinct chaperone-GR states *in vitro* as before using an MBP-tagged construct of GR (MBP-GR)^6,13,7^ and in the presence of CHIP. Specifically, to reconstitute the complex of Hsp70 with GR, we incubated MBP-GR with Hsp70 and the cochaperone Hsp40 in the presence of ATP. To reconstitute the GR-loading state, we incubated MBP-GR with Hsp70, Hsp40, Hop, and Hsp90 carrying the D93N mutation—which abrogates ATP binding and stalls the cycle at the Hsp90-GR-Hsp70-Hop intermediate—in the presence of ATP required for Hsp70. To reconstitute the GR-maturation state, we incubated MBP-GR with Hsp70, Hsp40, Hop, wild-type Hsp90, and p23 in the presence of ATP before addition of Bag-1—a co-chaperone of Hsp70 that blocks Hsp70-recapture of GR, thus enforcing directionality in the Hsp90/Hsp70 cycle and facilitating progression to the GR-maturation state^6,48^—as well as sodium molybdate, which stabilizes the post-ATP hydrolysis, closed conformation of Hsp90^49,50,12,7^. Previously, performing affinity purification for MBP-GR after assembling these different mixes *in vitro* has allowed us to discretely isolate functional Hsp70-GR, Hsp90-GR-Hsp70-Hop, and Hsp90-p23-GR complexes^6,7^. Finally, to test whether a chaperone intermediary is necessary for CHIP interaction with GR, we also incubated CHIP with MBP-GR on its own.

In line with the well-appreciated link between CHIP and Hsp70, affinity purification for MBP-GR from the mix of MBP-GR with Hsp70, Hsp40, and CHIP in the presence of ATP yielded a putative Hsp70-GR-CHIP complex, with all three components appearing in the pulldown eluate (**Fig. 1b, lane 2**). We next turned our attention to assessing Hsp90-GR-CHIP complexes, our primary mechanistic interest. Remarkably, affinity purification for MBP-GR from the GR-loading state mix plus CHIP co-purified not only Hsp90, Hsp70, and Hop, as to be expected, but also CHIP (**Fig. 1c, lane 2**), suggesting CHIP may be able to incorporate into the GR-loading state. In both the putative Hsp70-GR-CHIP and Hsp90-GR-Hsp70-Hop-CHIP complexes, GR is likely to be partially unfolded as in the corresponding parent complexes lacking CHIP. Speculating that CHIP might be sensitive to client foldedness, we hypothesized that CHIP would not be able to incorporate into the GR-maturation state, since GR is folded around ligand in this state^7^. To our surprise, affinity purification for MBP-GR from the GR-maturation state mix plus CHIP co-purified not only Hsp90 and p23, as to be expected, but also CHIP (**Fig. 1c, lane 3**), suggesting a putative four-component Hsp90-p23-GR-CHIP complex. Importantly, addition of CHIP to MBP-GR alone did not co-purify the E3 ligase (**Fig. 1c, lane 1**), indicating that CHIP does not recognize GR on its own in the absence of chaperone machinery.

CHIP co-purification with the GR-maturation state was unexpected and represents, to our knowledge, a first instance where a bona fide pro-folding Hsp90 state is observed to partner with an E3 ligase in a seemingly counterintuitive complex. To further validate the putative Hsp90-p23- GR-CHIP complex (**Fig. S1a**), we injected the pulldown eluate over a size-exclusion chromatography (SEC) column. Consistent with CHIP incorporation into the maturation complex, our SEC experiment showed a leftward shift in the elution peak relative to that for the Hsp90-p23- GR MBP-pulldown eluate (**Fig. 1d**). SDS-PAGE analysis of fractions from the peak corresponding to the CHIP-containing sample revealed the presence of all four components (**Fig. 1e**), strongly suggesting Hsp90-p23-GR-CHIP to be a bona fide complex. Glutaraldehyde crosslinking of the SEC fractions (**Fig. S1b**) revealed two major species at approximately the molecular weights expected for Hsp90-p23-GR and Hsp90-p23-GR-CHIP (**Fig. S1c**). Western-blot analysis of the crosslinked reactions showed positive signal against anti-Hsp90, anti-MBP, and anti-CHIP antibodies in the higher molecular-weight species (an anti-p23 Western was not performed) (**Fig. S1d**), supporting that it corresponded to Hsp90-p23-GR-CHIP.

### CHIP is functionally active in the Hsp90-p23-GR-CHIP complex

Having validated the Hsp90-p23-GR-CHIP complex, we sought to test whether it is functionally relevant—that is, whether it can mediate GR ubiquitylation. As mentioned, previously, we determined the cryo-EM structure of Hsp90-p23-GR and found that GR is folded around ligand^7^, consistent with *in vitro* measurements of GR’s ligand-binding ability in the presence of the full Hsp90/Hsp70 system^6^. In contrast, GR is partially unfolded in Hsp70-GR, with ligand-binding specifically inactivated^6^. It is thus possible that partially unfolded GR as presented by Hsp70 is in a more ready state for ubiquitylation, and that although the GR-maturation state can bind CHIP (e.g., with CHIP TPR domain binding to Hsp90 C-terminal EEVD), it is not productive for GR ubiquitylation. To test this hypothesis, we compared GR ubiquitylation by CHIP in these three different contexts: the receptor in the native state on its own, the partially unfolded receptor in complex with Hsp70, and the presumably native receptor as presented by the maturation complex. To allow for ubiquitylation activity, these different GR samples were briefly incubated with CHIP and a pre-formed E2∼ubiquitin (Ub) conjugate, UbcH5c∼Ub. The latter enabled us to bypass introduction of ATP into the ubiquitylation reactions, which would be required to activate ubiquitin were we to resort to an E1-E2-CHIP ubiquitylation cascade^51,52^ but which might induce turnover of the maturation state. To assess progress in GR ubiquitylation, we analyzed the reaction mixes with an anti-MBP Western (GR is MBP-tagged).

Strikingly, under our reaction conditions, we found that the GR-maturation state very efficiently promoted GR ubiquitylation by CHIP, with polyubiquitylated GR—that is, GR having received 4 or more ubiquitin molecules—readily detected at the highest CHIP concentration tested (**Fig. 2, lane 3**). Furthermore, across the CHIP titration tested, we consistently observe a greater amount of ubiquitylated GR overall in the GR maturation state compared with Hsp70-GR. Our results thus contradict the existing paradigm which focuses on Hsp70 as the preferred chaperone partner over Hsp90 for CHIP in coordinating client degradation^42^. Furthermore, they establish the Hsp90-p23-GR-CHIP complex to be a relevant state for entry into GR degradation. In contrast, in the absence of any chaperone machinery, CHIP was unable to ubiquitylate GR. This is consistent with the failure of GR alone to co-purify with CHIP (**Fig. 1c, lane 1**). Altogether, our findings demonstrate a strict requirement for chaperone presentation of GR to CHIP for ubiquitylation, with CHIP preferentially coordinating with Hsp90 over Hsp70 in mediating GR ubiquitylation.

**Figure 2:**
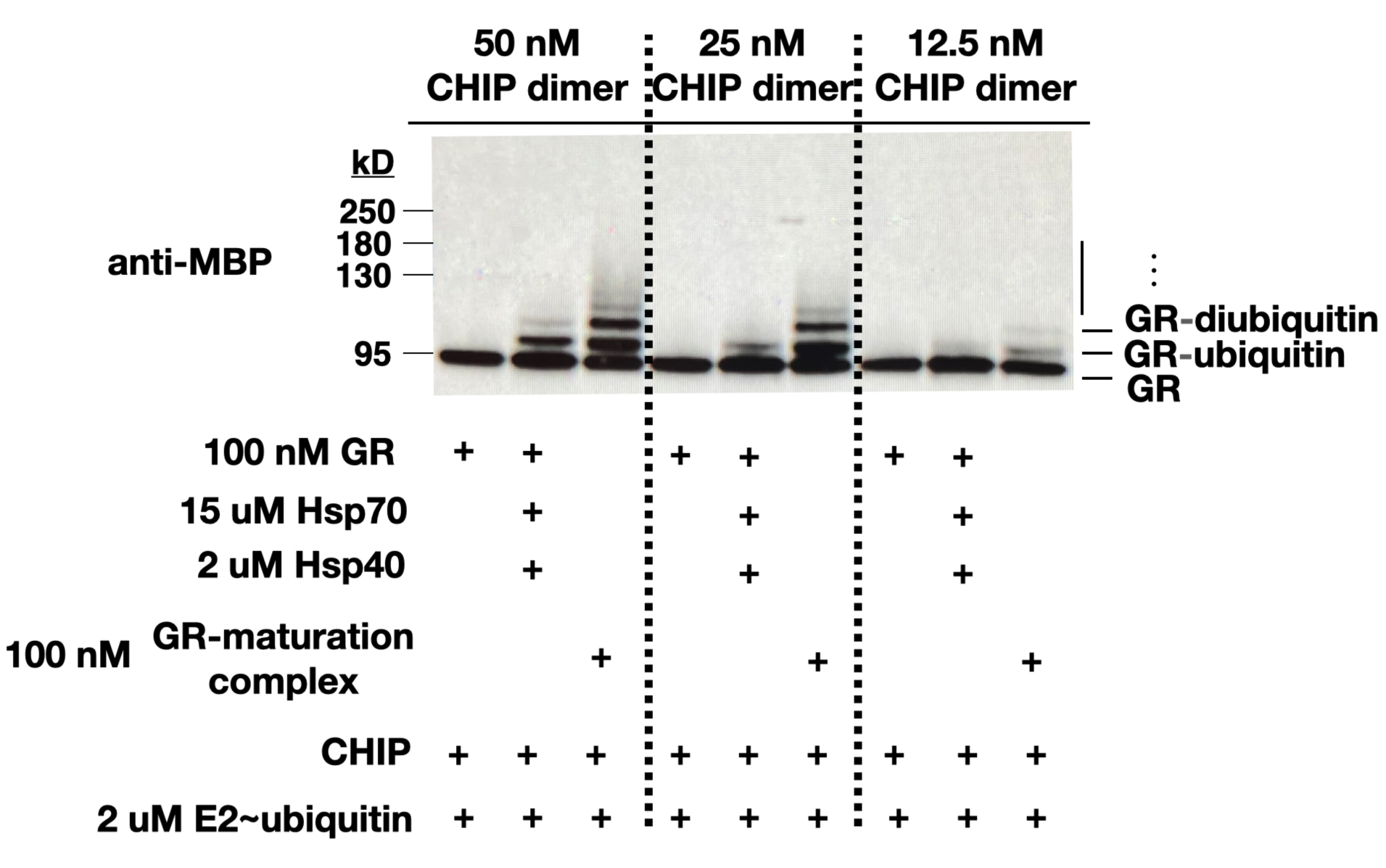
Hsp90-p23-GR-CHIP is a functionally active client-triage state. Western blot probed with MBP antibody against ubiquitylation reactions from incubating components as indicated. GR is N-terminally MBP-tagged.

### Hsp90-p23-GR-CHIP structure determination

Having shown Hsp90-p23-GR-CHIP represents a functionally active client-triage state, we attempted to determine the 3D architecture of the complex using single-particle cryo-EM. To prepare the sample for cryo-EM, MBP-GR was incubated with the complete Hsp90/Hsp70 system plus CHIP, and the maturation complex was enriched for with Bag-1 and stabilized with molybdate as before. Pulldown elution containing the four components—Hsp90, p23, GR, CHIP—was further stabilized by light glutaraldehyde crosslinking, and the crosslinked complex was purified by SEC (**Materials and Methods**, **Fig. S2a**). Deposition of the purified crosslinked sample onto functionalized carboxyl-PEG-5k (polyethylene glycol 5000) graphene oxide grids^53^ resulted in uniform particle distribution on the grids (**Fig. S2b**). The grids were used to collect a dataset on the Titan Krios.

A cryo-EM reconstruction at ∼14 Å was obtained resembling the GR-maturation complex, but bearing additional densities (**Figs. 3, S3 and S4**). Despite the low resolution, the general organization of the components in the complex can be assigned with high confidence. Hsp90 appears in the post-ATP hydrolysis, closed-dimer (Hsp90A/B) state bound to p23 and GR as in the previous maturation state structure^7^. Most strikingly, density presumably corresponding to CHIP emanates from the C-terminal domain (CTD) of Hsp90—a putative CHIP binding pose consistent both with the C-terminal EEVD motif on Hsp90 recruiting the TPR domain in CHIP, and with recent findings showing CTD binding to be a common motif for other TPR-containing cochaperones, FKBP51/52 and PP5^14,16^. Putative docking of a CHIP monomer suggests the TPR domain is oriented toward the base of Hsp90 and the U-box domain following to make putative contacts with the ligand-binding domain of GR as is presented by Hsp90 (**Fig. S5; Discussion**). Although CHIP has long been thought to be an obligate dimer since the report of its crystal structure^54^, emerging evidence suggests a dimer-monomer equilibrium in which the monomer state is also catalytically competent^55^.

**Figure 3:**
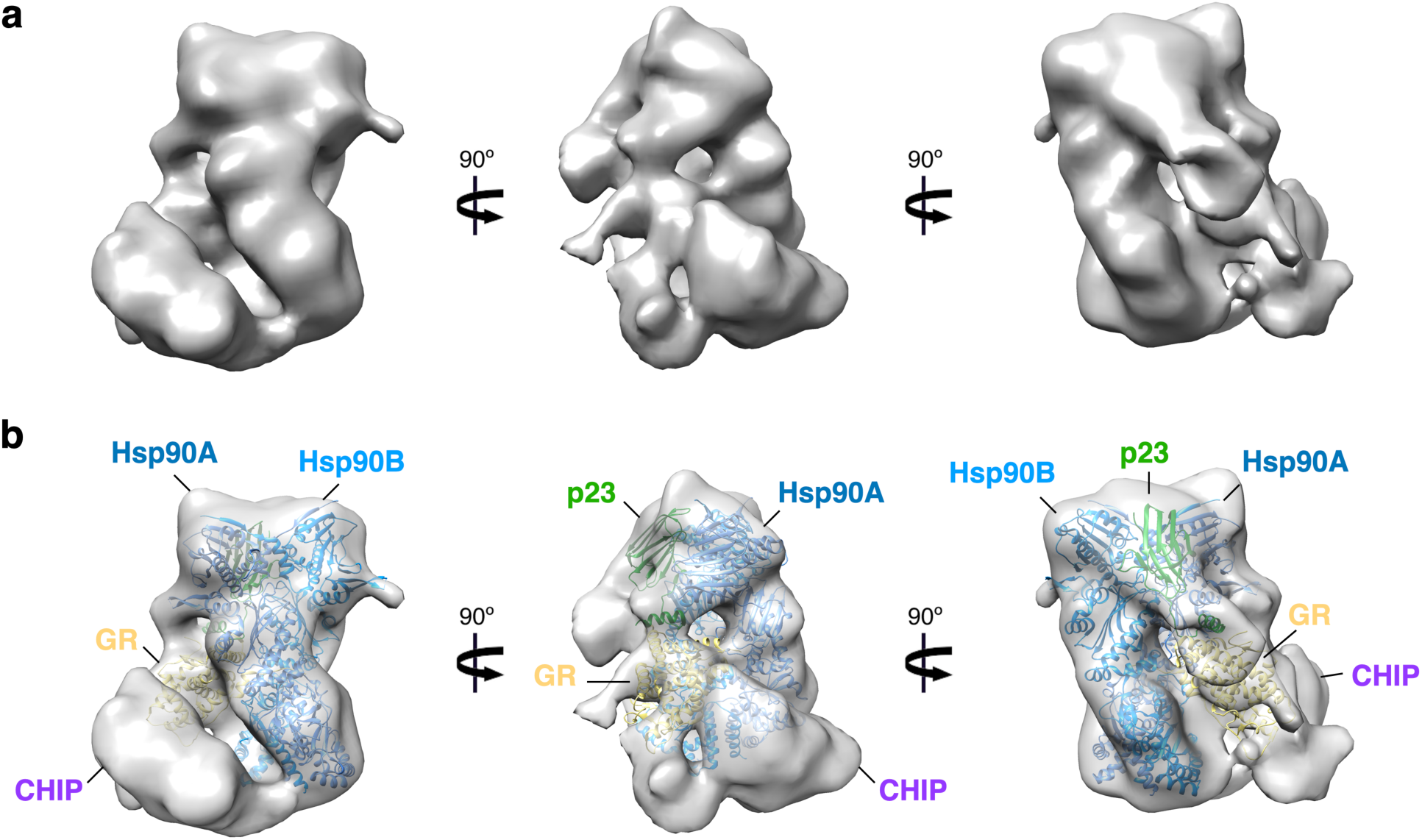
Architecture of Hsp90-p23-GR-CHIP. **a**, Cryo-EM reconstruction of Hsp90-p23-GR-CHIP. **b**, Cryo-EM reconstruction as in (**a**) with placement of the GR-maturation complex (PDB: 7KRJ), in which the Hsp90 dimer—composed of protomers Hsp90A and Hsp90B—is colored blue, p23 is colored green, and GR is colored yellow. Extra density likely corresponding to CHIP is highlighted. See also **Fig. S5**.

Focused classification with masks and subtraction failed to show clearer CHIP density, suggesting that additional stabilization and/or a far larger data set would be required. To more properly assess potential changes to the GR fold in our sample, we used a mask to guide focused classification on GR (**Fig. S3**), yielding one class containing clear GR density. Upon reconstructing this subset of particles in RELION, we observed partial p23 occupancy. As the presence or absence of p23 could have consequences on the GR fold, given that we previously observed a helix tail on p23 to directly stabilize GR in the Hsp90-p23-GR complex^7^, we further performed focused classification for p23, yielding one class containing well-resolved p23 density. Global refinement on this class of particles yielded a map of 3.53 Å resolution (**Figs. 4, S3, S6**).

**Figure 4:**
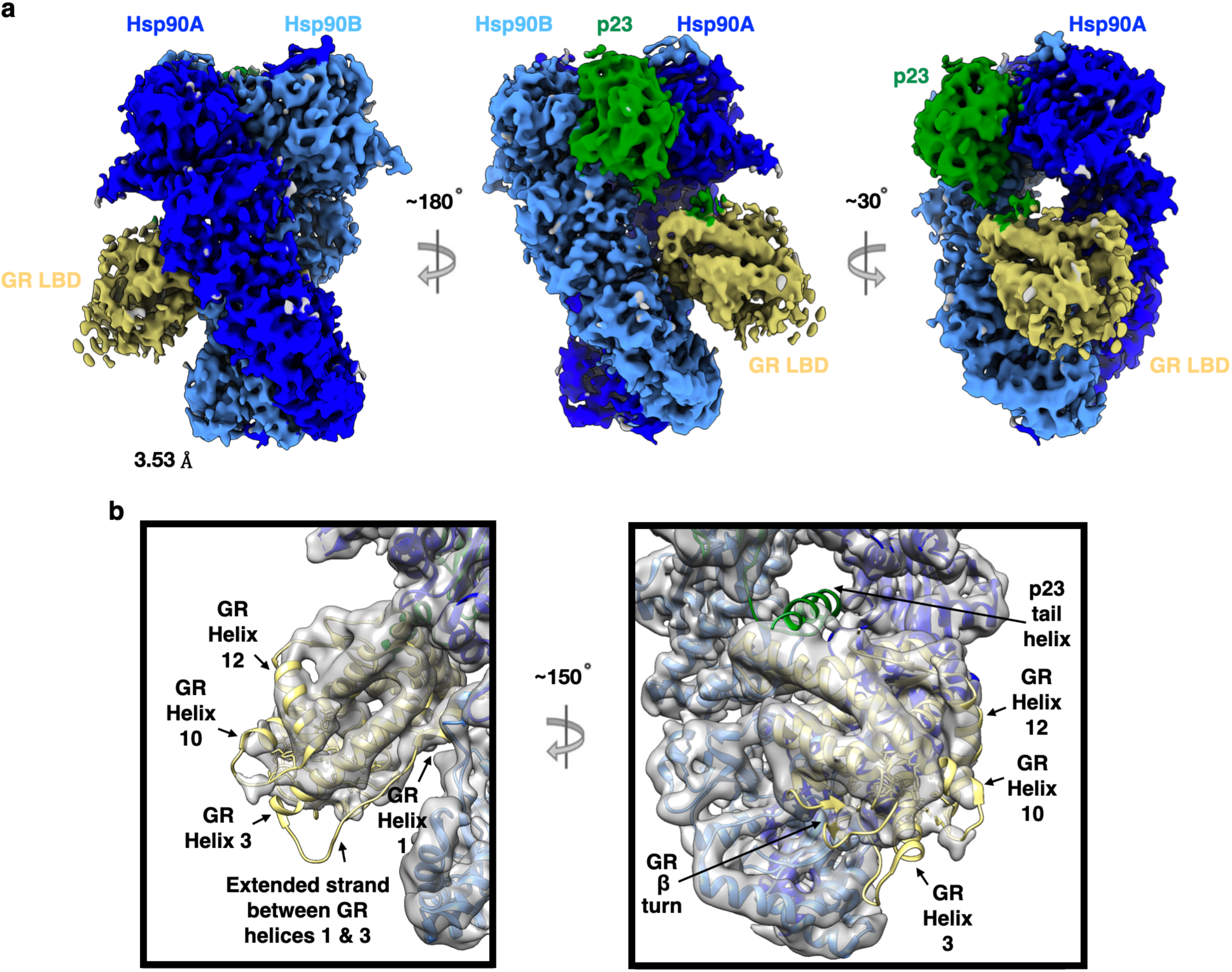
GR is ligand-free and partially disordered. **a**, Cryo-EM reconstruction of an apo Hsp90-p23-GR complex, showing GR is ligand-free and partially disordered. Hsp90A, dark blue; Hsp90B, light blue; GR, yellow; p23, green. Color scheme is maintained throughout. **b**, The GR ligand-binding domain in (**a**) is shown in detail. The previously determined GR-maturation complex (PDB: 7KRJ^7^) is fitted into the cryo-EM reconstruction in (**a**) after applying a Gaussian filter at 1.314 to the map and thresholding the resulting reconstruction at 0.183. In the PDB structure, Hsp90 is colored blue, p23 is colored green, and GR is colored yellow. Substructures of interest of the GR ligand-binding domain as well as the p23_tail-helix_ are highlighted. Notably, our cryo-EM map shows density is missing for the extended strand between GR_Helix 1_ and GR_Helix 3_, the first two helical turns of GR_Helix 3_, the β-turn, and the final helical turn of GR_Helix 10_. Moreover, p23_tail-helix_ is also poorly resolved.

Overall, we observed no changes to GR position or orientation relative to the previously determined GR-maturation state structure^7^. Furthermore, the core of the so-called ‘helical sandwich’^56,57^ in the top half of the ligand-binding domain—comprised of a top layer of helices 1 and 3, a middle layer of helices 4, 5, 8, and 9, and a bottom layer of helices 7 and 10—is well-assembled, as in the ligand-bound state. In contrast with the GR-maturation state structure, however, we do not observe density for ligand, even though ligand was present during the chaperone incubation during sample preparation. Equally surprisingly, density is clearly missing or disordered for the extended strand between GR_Helix 1_ and GR_Helix 3_ as well as features that comprise the distal portion of the ligand-binding pocket (**Fig. 4b**). These features include the first two helical turns of GR_Helix 3_ (which notably contain M560, normally engaged in van der Waals contact with ligand), the β-turn (containing F623, which normally engages in van der Waals contact with ligand), and the final helical turn of GR_Helix 10_ (containing T739, which normally engages in hydrogen bonding with ligand) (refer to **Fig. 4b** and **Fig. S7** to see how contour level affects the resolution of these features). Remarkably, however, despite these structural differences and the absence of ligand, GR_Helix 12_ still adopts an agonist-bound position, in agreement with several ligand-free nuclear receptor structures also showing helix 12 in this ‘active’ position^58,59,60,61^. Furthermore, the conserved β-sheet comprised of the two C-terminal β strands in GR appears intact. In contrast, the so-called p23_tail-helix_—found previously to dock onto a hydrophobic patch on GR in a stabilizing interaction^7^—is largely missing, suggesting it might dynamically undock. Although it is unclear to us how this would happen, given the extensive contacts normally observed between the two interaction partners, our previous biochemistry experiments had shown that the p23_tail-helix_ potentiates ligand-binding by GR^7^, and thus its poor resolution in our map would seem to be consistent with the apo status of GR in our dataset. These details aside, overall, our results suggest CHIP favors or causes a partially disordered GR as substrate.

## Discussion

Taking advantage of our unique ability to reconstitute *in vitro* key folding intermediates along the Hsp90 cycle for the glucocorticoid receptor (GR)^6,13,7,14^, we find that CHIP recognition of GR is strictly dependent on chaperone presentation and also show that both Hsp90-GR loading and maturation complexes can bind CHIP. While the GR-maturation state was previously thought by us to be a strictly pro-folding state^7^, our present work revises this and reveals that in the presence of CHIP, the GR-maturation state in fact promotes efficient ubiquitylation of partially unfolded GR, in this way functioning as a client-triage state, facilitating both folding and degradation outcomes (**Fig. 5**).

**Figure 5:**
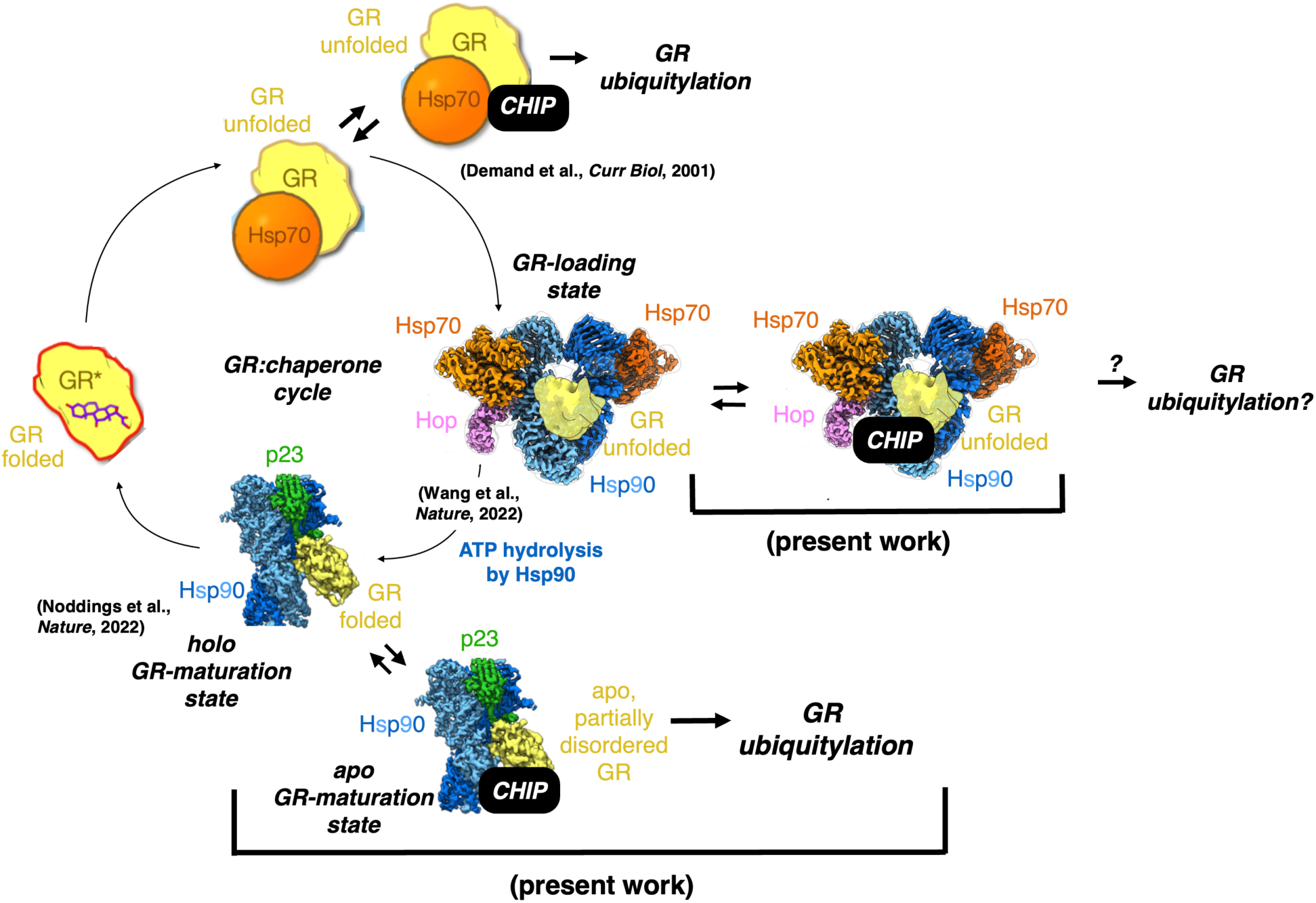
Model of Hsp90-dependent triage of GR via CHIP. The GR:chaperone cycle shown in Fig. 1a is elaborated with exit points to chaperone- dependent CHIP-containing states discussed or identified in the present work. Based on results reported herein, we propose ubiquitylation as another basal outcome (alongside folding) of GR’s progression along the Hsp90 cycle, via CHIP intercepting an apo GR-maturation state and/or potentially CHIP intercepting the GR-loading state as well. In this way, Hsp90 actively mediates protein triage decisions in response to client foldedness.

In contradiction with an existing paradigm highlighting Hsp70 as the preferred chaperone partner for CHIP in coordinating client degradation^42^, we find that CHIP preferentially ubiquitylates GR when presented in the Hsp90 maturation state relative to GR as presented by Hsp70 alone. Observed differences in GR ubiquitylation efficiencies may stem from a combination of differences in (i) the binding kinetics underlying CHIP engagement with the different chaperone-GR complexes; (ii) the processivity of ubiquitylation in the different chaperone-GR-CHIP complexes; and (iii) the ubiquitylation sites on GR and their accessibility in the different chaperone-GR-CHIP complexes. Quantitative analyses in the future of these various aspects of the different complexes and the corresponding ubiquitylation reactions should elucidate how two different chaperones can coordinate with a common E3 ligase to ubiquitylate the same client substrate.

Based on the observation that CHIP can ubiquitylate the Hsp90 client p53 on its own, with chaperones only enhancing this activity^42,47^, one might have expected a similar behavior with GR. In contrast, our pulldown and ubiquitylation assays show instead that GR has a strict dependence on chaperone presentation for CHIP engagement. This is in line with other studies also demonstrating a strict chaperone (generally Hsp70) requirement for CHIP activity^46,44,45^, perhaps because GR and those clients are more fully folded than p53. Chaperones would thus seem to play an active role in guiding CHIP engagement with client/GR, perhaps by remodeling client/GR to a CHIP-sensitive conformational state, in addition to providing additional affinity by binding the chaperones via their flexible C-terminal EEVD tails.

In agreement with the hypothesis of Hsp90 presenting GR in a CHIP-sensitive conformational state that is distinct from the native state, our cryo-EM analysis reveals a first ligand-free and partially disordered GR structure bound to the post-ATP hydrolysis, closed Hsp90 dimer, likely existing in equilibrium with ligand-bound GR as previously observed^7^. Although the core of the ‘helical sandwich’ is well-assembled and GR_Helix 12_ notably still adopts an agonist-bound position, upon setting our map to a moderate contour level, we find that density is missing for (i) ligand, (ii) GR substructures comprising the distal end of the ligand-binding pocket, as well as (iii) the extended strand between GR_Helix 1_ and GR_Helix 3._ The likely disordered nature of this extended strand between GR_Helix 1_ and GR_Helix 3_ is especially intriguing in light of the ligand’s absence in our map. It would appear to corroborate previous single-molecule force spectroscopy experiments on GR demonstrating that hormone binding into the ligand-binding pocket is gated by the opening or closing of the first 33 amino acids of the ligand-binding domain^62^—which encompass GR_Helix 1_ and precisely this extended strand between helices 1 and 3. In the GR-maturation state structure, GR_Helix 1_ packs up against an amphipathic helical hairpin in the C-terminal domain of Hsp90 in a stable interaction, and we had expected this to remain the case in our current structure (**Fig. 4b**) (although a high-resolution analysis would be required to confirm this). Our structural results instead implicate the order-disorder transition of the extended strand between GR_Helix 1_ and GR_Helix 3_ to potentially be the more relevant determinant gating ligand entry into and dissociation from the ligand-binding pocket in the GR-maturation state, and this order-disorder transition may in turn be coupled to the folding of the β-turn as well as segments of helices 3 and 10 in GR that are missing in our reconstruction.

The speculative nature of this hypothesis notwithstanding, our finding of a partially disordered GR state in our cryo-EM dataset would suggest that CHIP can partially disorder or prefers partially disordered GR as substrate, which could explain why our pulldown and ubiquitylation assays did not reveal an interaction between CHIP and native-state GR alone. Further analysis of the Hsp90-p23-GR-CHIP complex would be needed to elucidate how CHIP might recognize and/or enrich for partially disordered GR. Until then, our present results are already noteworthy for demonstrating for the first time that a partially unfolded GR intermediate can still be presented by a post-ATP hydrolysis Hsp90 state, contrary to structures available to date^7,14^. In other words, GR folding is not the sole, deterministic outcome following ATP hydrolysis by Hsp90. Instead, we find that CHIP can incorporate into post-hydrolysis Hsp90-GR and efficiently mediate GR ubiquitylation. Thus, we have shown that the post-hydrolysis Hsp90-GR state is an instance along the Hsp90 cycle where client foldedness could be utilized by Hsp90 as a metric for targeting damaged GR for degradation—that is, GR that does not properly fold around ligand fast enough might preferentially be ubiquitylated by CHIP and as such triaged for degradation (**Fig. 5**).

Our low-resolution reconstruction of Hsp90-p23-GR-CHIP additionally suggests CHIP might also undergo conformational changes from the dimer structure previously captured in crystallography experiments^54^, as the extra density in our map would accommodate only one CHIP monomer. Previously, the crystal structure of CHIP revealed an asymmetric dimer, in which the protein adopts two different folds in the two different protomers. For one, the long helical hairpin (HH) domain is folded differently in the two different protomers. More importantly, one protomer (CHIP_protomer 1_; chains A and C in PDB 2C2L) is catalytically incompetent, as it features an inhibitory interaction between the TPR domain and the U-box domain that would prevent the U-box domain from engaging with an E2 enzyme (**Fig. S5a**), while the other protomer (CHIP_protomer 2_; chains B and D in PDB 2C2L) does not and would thus be catalytically competent (**Fig. S5b**). Using the “Fit in Map” function in Chimera, we were able to dock both protomer structures into the extra density (**Fig. S5c** and **Fig. S5d, respectively**). In both cases, the TPR domain was best positioned at the C-terminal domain of Hsp90, from where it can present the U-box domain to the GR client, while the HH domain would appear to protrude from the map unresolved. Admittedly, at this resolution the docking in both cases is highly speculative. However, it is intriguing to note that based on the docking of CHIP_protomer 2_—i.e., the catalytically active protomer—and then sequentially aligning the available crystal structures of (1) the U-box domain in complex with the E2 enzyme UbcH5a (PDB 2OXQ) and (2) the E2 enzyme Ubch5b in complex with ubiquitin (PDB 3A33), we were able to generate a three-dimensional model of CHIP-GR-p23-Hsp90 in complex with an E2∼ubiquitin conjugate (**Fig. S8a**). Importantly, the ubiquitin molecule presented in this way would sterically clash with folded GR as is resolved in the GR-maturation complex (**Fig. S8b**) but would potentially be compatible with the partially unfolded GR intermediate observed here. Much higher resolution structural information is clearly desired to elucidate how CHIP recognizes GR and facilitates its ubiquitylation.

More broadly speaking, we expect that multiple client-triage states exist and remain to be characterized. For example, we have found that CHIP also co-purifies with the GR-loading state— suggesting the GR-loading state to be yet another client-triage state, although biochemical characterization of GR’s susceptibility to ubiquitylation in this state awaits. Even so, our present work on Hsp90-p23-GR-CHIP sufficiently calls for the concept of the Hsp90-cycle to be updated. Rather than facilitating folding as a singular outcome, we have demonstrated that a folding state can similarly support ubiquitylation in the presence of an E3 ligase. In other words, upon association with Hsp90, a client protein could be triaged for either a folding outcome or degradation, presumably as part of a protein quality control mechanism (**Fig. 5**). We propose that the net balance of client folding vs. degradation along the Hsp90 cycle is dictated by a balance of the lifetime of the folding states and the kinetics of E3 ligase/deubiquitinase binding and activities. Previously, in having delineated the requirement for an unfolding step (as mediated by Hsp70) in the Hsp90-folding cycle for GR, we conjectured that such a tight coupling between folding and unfolding provides opportunities for regulatory control and to coordinate signaling with protein homeostasis^6^. We expect these same opportunities are afforded by the even tighter coordination between folding and degradation that we have begun to reveal here, where a single Hsp90-client state, depending on the cochaperone factors with which it partners, has the potential to guide a client protein towards two opposing fates.

## Acknowledgments

We thank members of the Agard Lab, past and present, for helpful discussions. We thank David Bulkley, Glenn Gilbert, Zanlin Yu, and Eric Tse from the W. M. Keck Foundation Advanced Microscopy Laboratory at the University of California, San Francisco (UCSF) for EM facility maintenance and training. We thank Matt Harrington and Joshua Baker-LePain for computational support with the UCSF Wynton cluster. We thank Daniel Asarnow for valuable comments on the manuscript. We thank Yu Chen for help with EMDB deposition.

## Author Contributions

C.M.C. designed and executed biochemical experiments, cryo-EM sample preparation, data collection, and data processing, and wrote the initial manuscript draft. F.W. generated PEG-5k carboxylate affinity grids and provided grid blotting conditions. C.M.N. provided protocols related to GR expression and purification and GR pull-down assays; the GR-DBD LBD protein used in Fig. 2; as well as valuable comments on the manuscript. C.M.C. and D.A.A. conceived the project, interpreted the results, and revised the manuscript.

---

**Figure S1:**
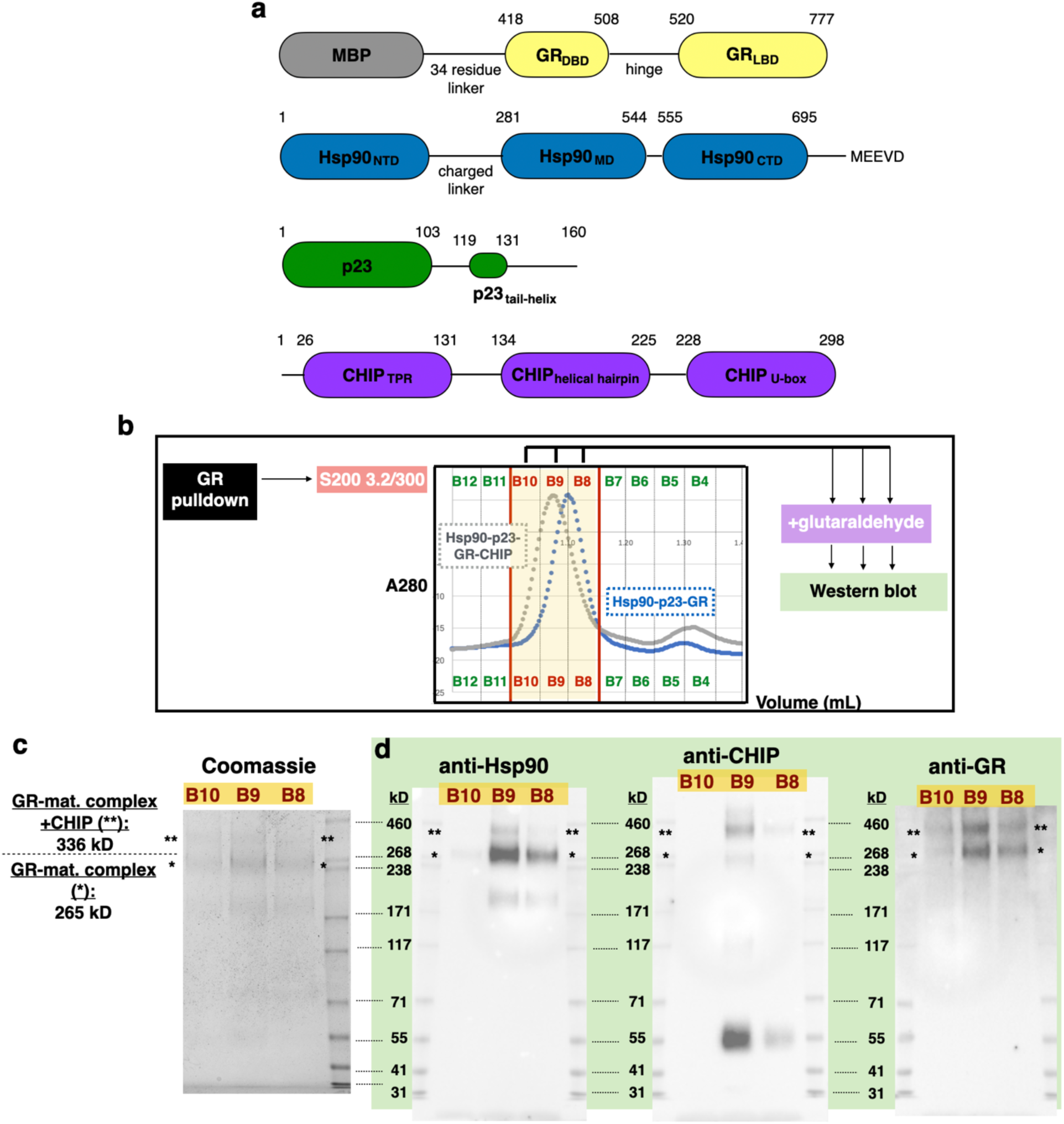
Crosslinking characterization of Hsp90-p23-GR-CHIP. **a**, Domain organization of the proteins in the Hsp90-p23-GR-CHIP complex. **b**, Schematic of workflow leading to the crosslinking products shown in (**c**) and (**d**). MBP-GR pulldown of Hsp90- p23-GR-CHIP was injected over S200 3.2/300. Fractions B10, B9, and B8 from gel filtration containing Hsp90, p23, GR, CHIP (as shown in Fig. 1e) were then treated with 0.05% glutaraldehyde and then visualized by SDS-PAGE in (**c**) and Western blot in (**d**). **c**, Coomassie-stained SDS-PAGE analysis of crosslinking reactions as described in (**b**). Single asterisk (*) denotes gel band approximating the molecular weight of GR-maturation complex alone (265 kD). Double asterisk (**) denotes gel band approximating the molecular weight of Hsp90-p23- GR-CHIP (336 kD). **d**, Western blots probed with Hsp90, CHIP, and GR antibodies against crosslinking reactions as described in (**b**). Single and double asterisks (*, **) denote entities as in (**b**).

**Figure S2:**
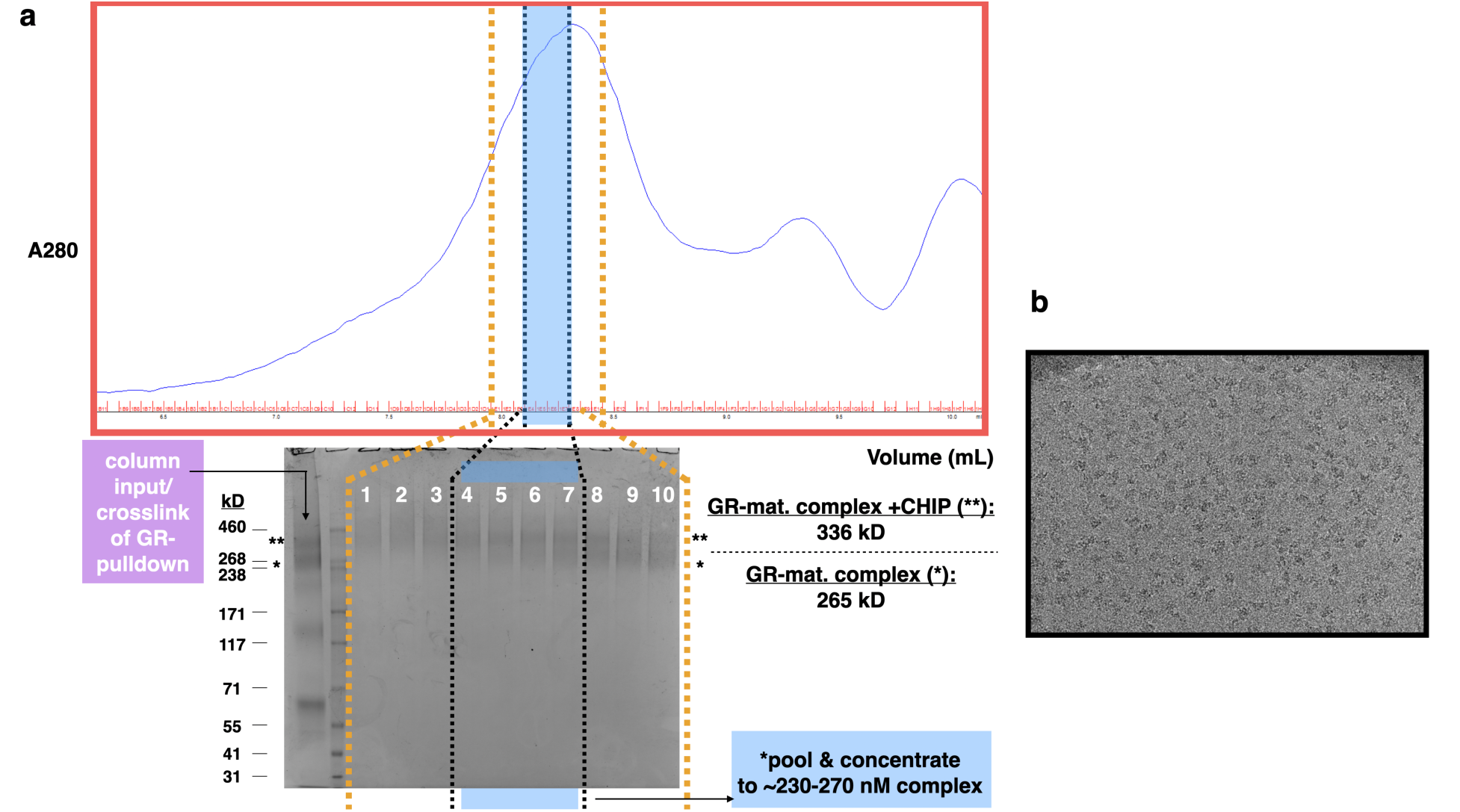
Sample Purification for cryo-EM. **a***, Top*, Shodex KW-804 gel-filtration profile of the glutaraldehyde-crosslinked MBP-GR pulldown of Hsp90-p23-GR-CHIP. *Bottom*, Coomassie-stained SDS-PAGE analysis of column input as well as gel-filtration fractions. Single asterisk (*) denotes gel band approximating the molecular weight of GR-maturation complex alone. Double asterisk (**) denotes gel band approximating the molecular weight of Hsp90-p23-GR-CHIP. Gel-filtration fractions denoted with a horizontal blue band were pooled then concentrated for vitrification. **b**, Representative electron micrograph for the cryo-EM dataset.

**Figure S3:**
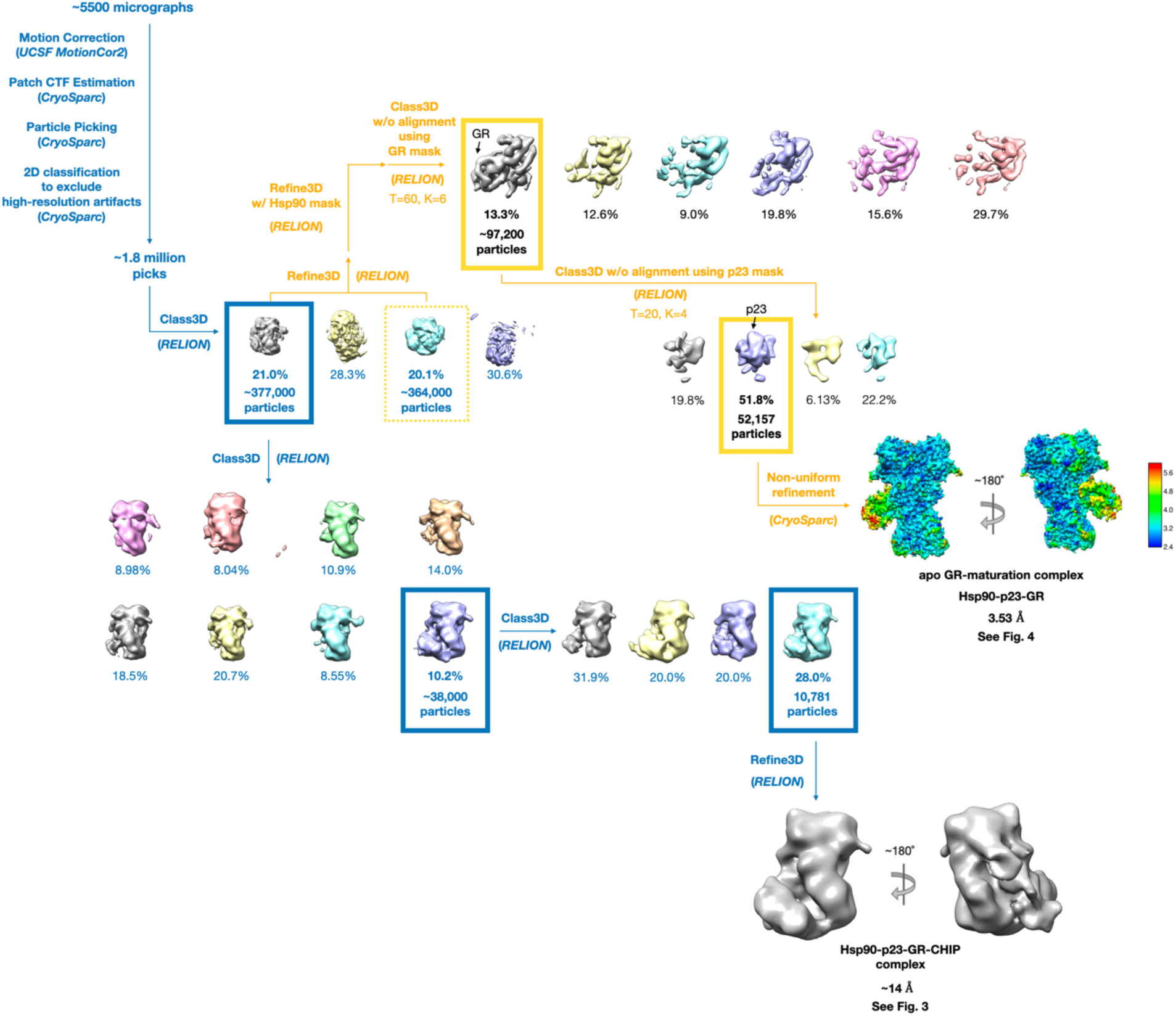
Cryo-EM single-particle imaging processing pipeline of the Hsp90-p23-GR-CHIP and apo Hsp90-p23-GR complexes. Cryo-EM data processing procedure yielding the low-resolution reconstruction of the Hsp90-p23- GR-CHIP complex shown in Fig. 3 as well as the high-resolution analysis of the apo Hsp90-p23- GR complex shown in Fig. 4. Processing steps leading to the former are colored in blue, while divergent processing steps leading to the latter are colored in orange. In both cases, selected particle subsets are highlighted with a colored box.

**Figure S4:**
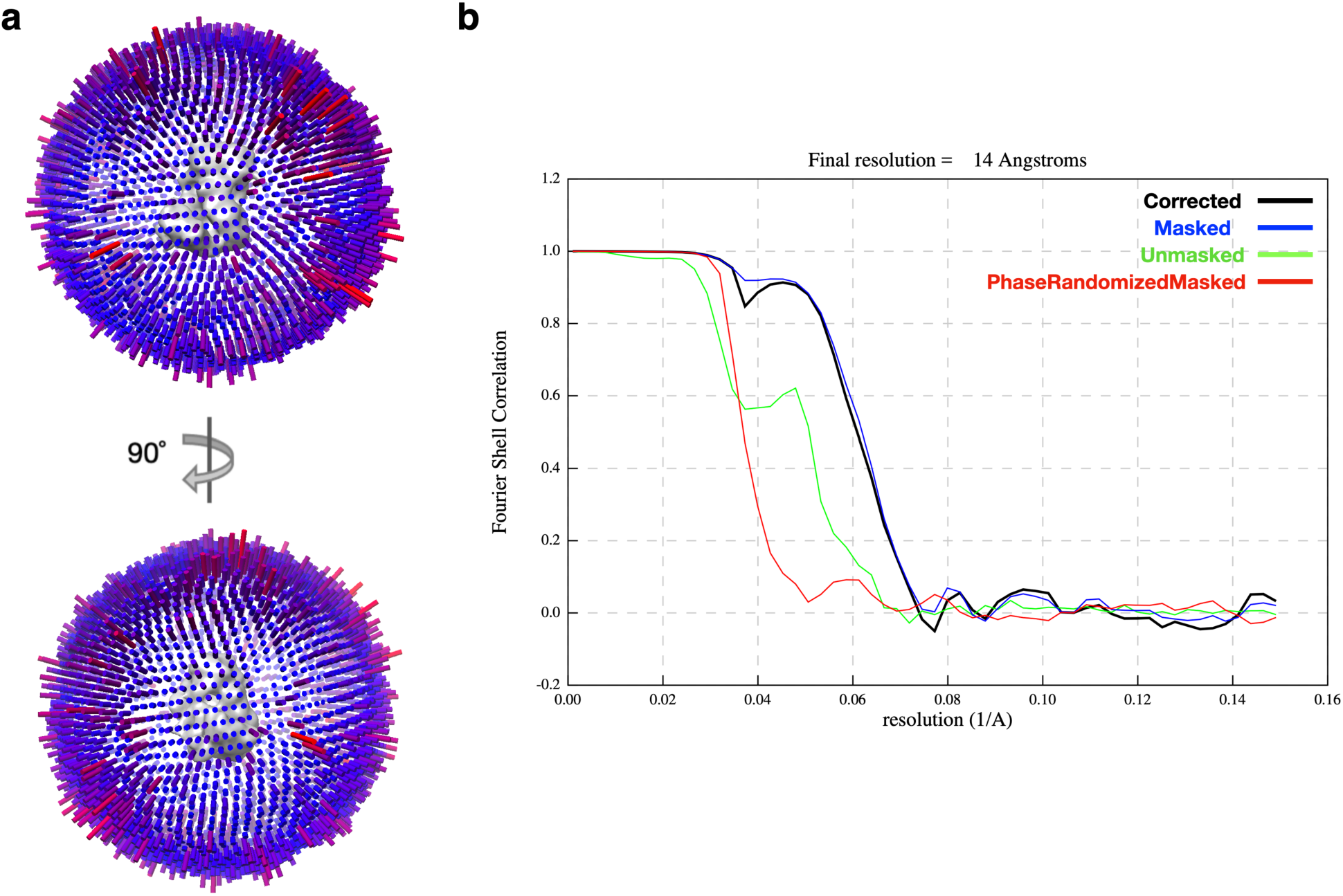
Cryo-EM single-particle analysis of Hsp90-p23-GR-CHIP complex. **a**, Euler angle distribution in the final reconstruction shown in Fig. 3a. Orthogonal views of the reconstruction are shown with front view (top) and side view (bottom). **b**, Gold Standard FSC for the global cryo-EM reconstruction.

**Figure S5:**
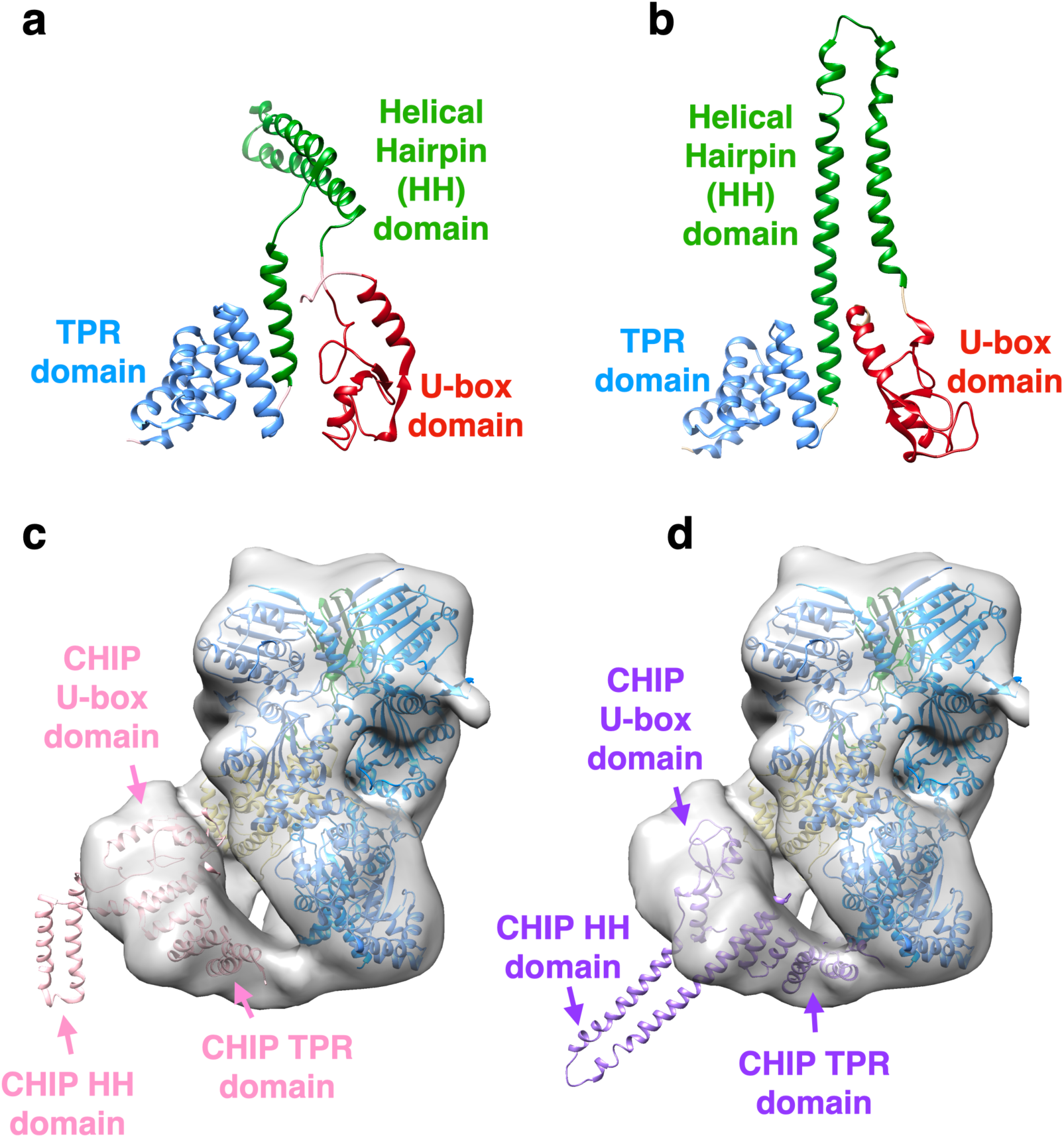
Docking of CHIP into the Hsp90-p23-GR-CHIP EM density. **a**, Structure of CHIP_protomer 1_, the catalytically incompetent protomer of CHIP (PDB: 2C2L, chain A/C), with domains identified. **b**, Structure of CHIP_protomer 2_, the catalytically competent protomer of CHIP (PDB: 2C2L, chain B/D), with domains identified. **c**, Cryo-EM reconstruction of Hsp90- p23-GR-CHIP with placements of the GR-maturation complex (PDB: 7KRJ) as in Fig. 3b as well as CHIP_protomer 1_ from (**a**) colored in pink. **d**, Cryo-EM reconstruction of Hsp90-p23-GR-CHIP as in Fig. 3b with placements of the GR-maturation complex (PDB: 7KRJ) as in Fig. 3b as well as CHIP_protomer 2_ from (**b**) colored in purple.

**Figure S6:**
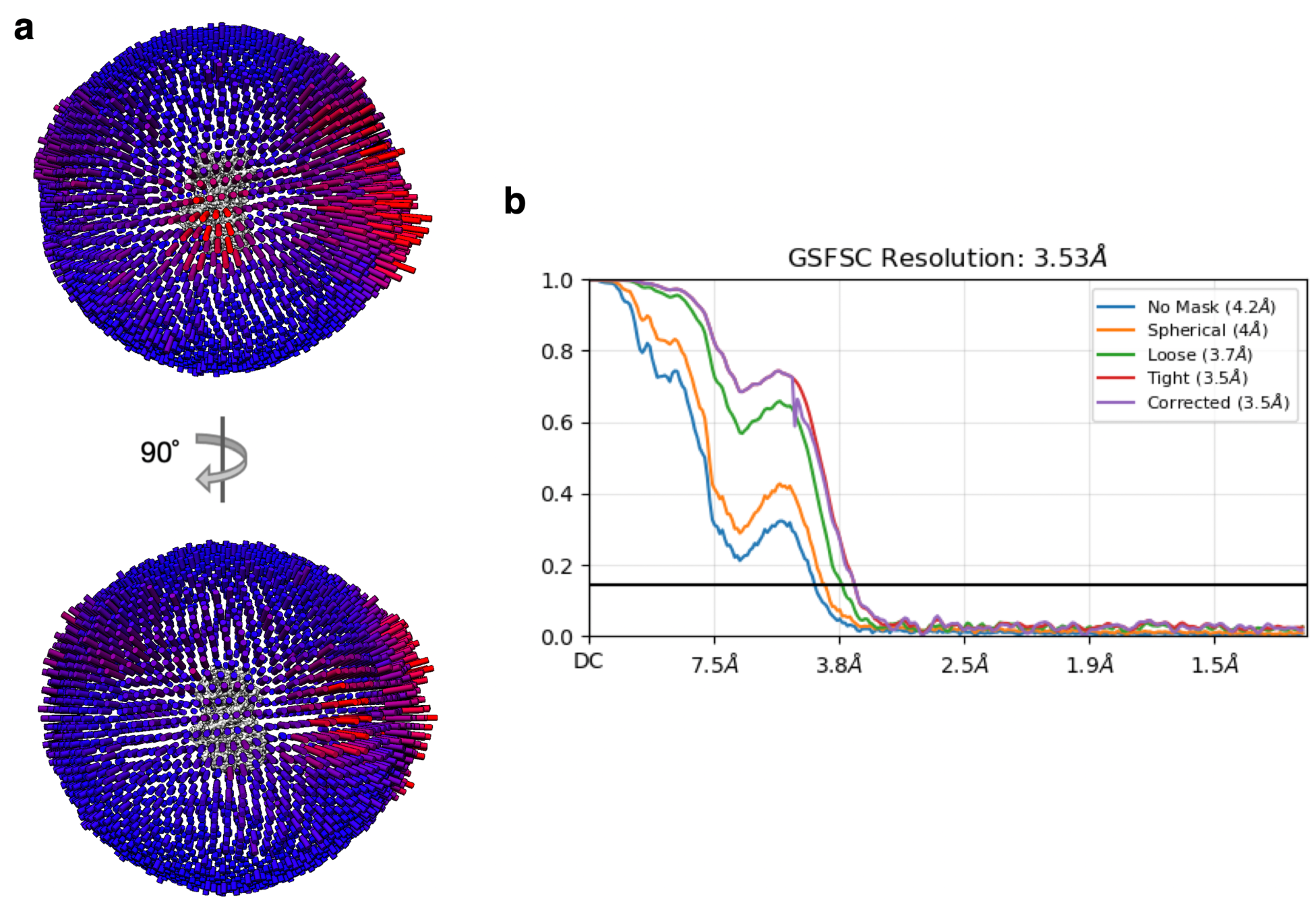
Cryo-EM single-particle analysis of the apo Hsp90-p23-GR complex. **a**, Euler angle distribution in the final reconstruction shown in Fig. 4a. Orthogonal views of the reconstruction are shown with front view (top) and side view (bottom). **b**, Gold Standard FSC for the global cryo-EM reconstruction.

**Figure S7:**
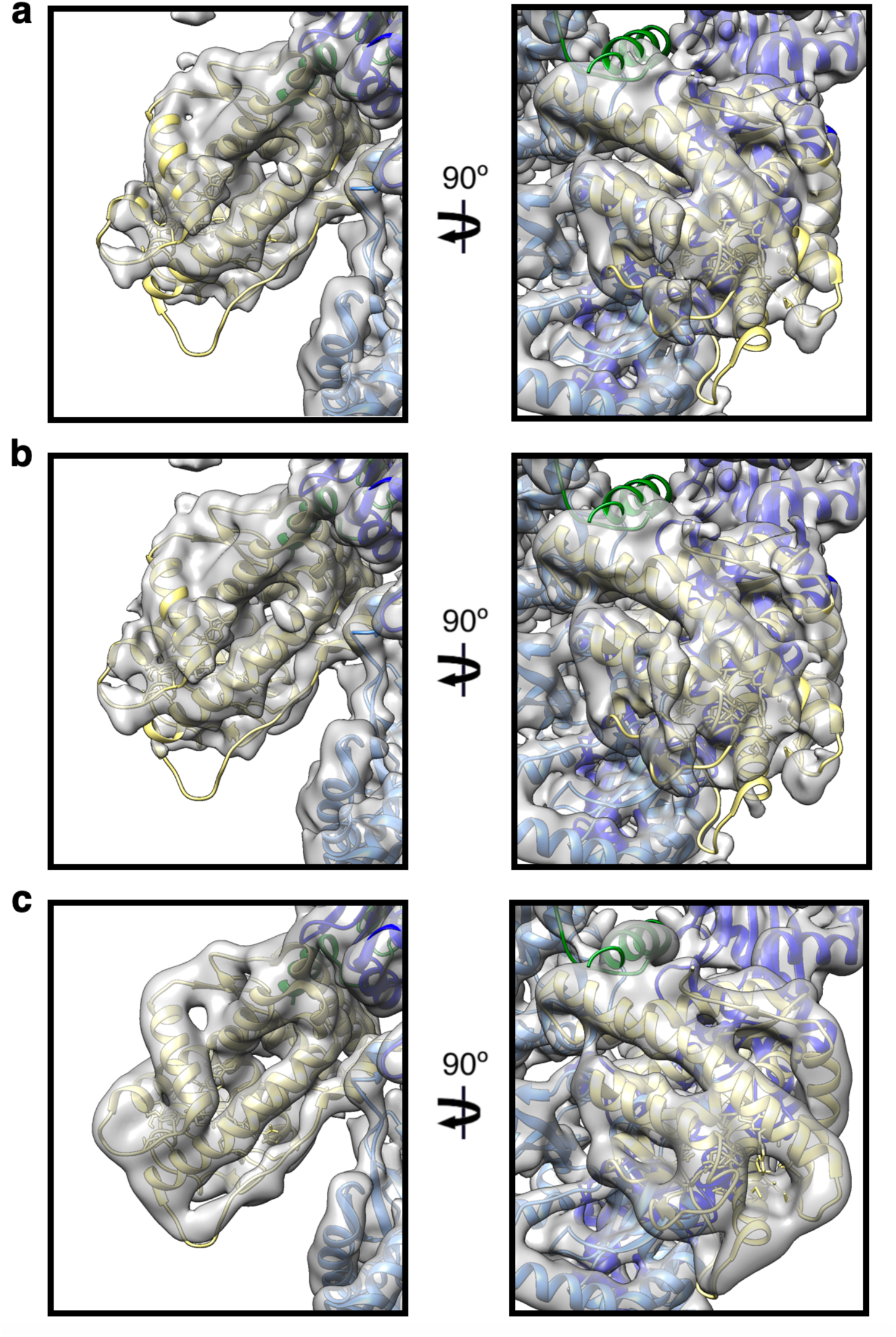
Varying degrees of GR ligand-binding domain disorder are suggested at varying contour levels. **a**, Visualization of the cryo-EM reconstruction in Fig. 4a after applying a Gaussian filter at 1.314 and thresholding at 0.173. **b**, Visualization of the cryo-EM reconstruction in Fig. 4a after applying a Gaussian filter at 1.314 and thresholding at 0.163. **c**, Visualization of of EMD-23004 (GR-maturation complex as reported in Noddings et al., 2022) after applying a Gaussian filter at 1.67 and thresholding at 0.0104. Unlike in the above two figures and Fig. 4b, density is readily apparent for the entire GR ligand-binding domain and p23_tail-helix_.

**Figure S8:**
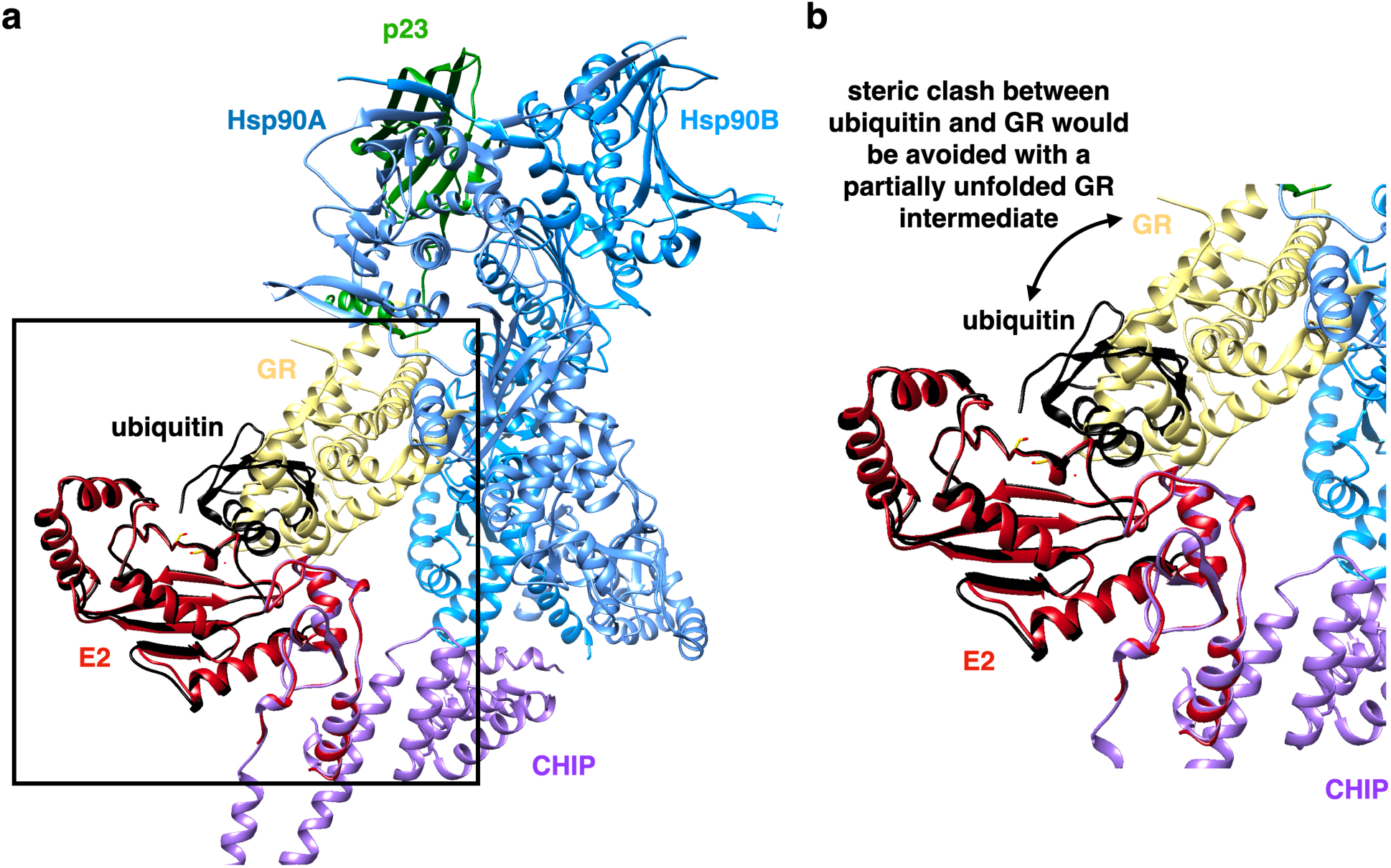
Model of Hsp90-p23-GR-CHIP in complex with an E2∼ubiquitin conjugate. **a**, Following fitting of the GR-maturation complex (PDB: 7KRJ) and CHIP_protomer 2_ (PDB: 2C2L, chains B/D) into the Hsp90-p23-GR-CHIP EM density map as in **Fig. S5d**, the structure of the U-box domain in complex with the E2 enzyme UbcH5a (PDB: 2OXQ; colored in red) was first aligned to CHIP. Next, the structure of the E2 enzyme Ubch5b in complex with ubiquitin (PDB: 3A33; colored in black) was aligned to 2OXQ. Box highlights the zoomed-in view presented in (**b**). **b**, Zoomed in view of (**a**) showing ubiquitin would sterically clash with folded GR as resolved in the GR-maturation complex. It is possible a partially unfolded GR intermediate might be accommodated on the other hand.

**Supplementary Data Table 1.** Cryo-EM data collection and processing.

|  | Hsp90-p23-GR-CHIP<br>(EMDB-?????) | apo Hsp90-p23-GR<br>(EMDB-?????) |
| --- | --- | --- |
| Magnification | 130,000 | 130,000 |
| Voltage (kV) | 300 | 300 |
| Electron exposure (e-/Å<sup>2</sup>) | 66 | 66 |
| Defocus range (μm) | 0.8-2.0 | 0.8-2.0 |
| Pixel size (Å) | 0.652 | 0.652 |
| Symmetry imposed | C1 | C1 |
| Initial particle images (no.) | ~1,800,000 | ~1,800,000 |
| Final particle images (no.) | 10,781 | 52,157 |
| Map resolution (Å) | 14.4 | 3.53 |
| FSC threshold | 0.143 | 0.143 |

## Methods

### Protein Expression and Purification

Human Hsp90a, Hop, Hsp70, Hsp40, p23, Bag1, and CHIP were separately expressed as constructs with an N-terminal 6xHis-TEV-cleavage tag in BL21RIL. Cells were grown in TB at 30 degC or 37 degC until OD600 reached 0.4, at which point the temperature was lowered to 16 degC (18 degC for CHIP). When the OD600 reached 0.8, IPTG was added to final concentrations of 500 uM for Hsp90a and 1 mM for all other proteins for overnight induction. Cells were harvested and lysed by an EmulsiFlex-C3 (Avestin) in 50 mM Tris pH 8, 750 mM KCl, 10% glycerol, 10 mM imidazole, 6 mM b-mercaptoethanol, and protease inhibitor pills (Roche). The lysate was clarified by centrifugation and the soluble fraction was affinity purified by gravity column with Ni-NTA affinity resin (QIAGEN). The protein was eluted with 30 mM Tris pH 8, 150 mM KCl, 400 mM imidazole, and 6 mM b-mercaptoethanol. For Hsp90, an extra wash step with 0.1% Tween20 and 2 mM ATP/MgCl2 was added to the Ni-NTA resin before eluting. The 6x-His tag was removed with TEV protease during overnight dialysis in 30 mM Tris pH 8, 100 mM KCl, and 6 mM bME. Cleaved protein was then loaded on a MonoQ 10/100 GL (GE Healthcare) ion exchange column and eluted with a linear gradient of 100mM-500mM KCl for Hsp90 and 30 mM-500mM KCl for all others. Protein was further purified by size exclusion with a Superdex S200 16/60 (GE Healthcare) or Superdex S75 16/60 (GE Healthcare) in either 30 mM HEPES pH 7.5, 50 mM KCl, 10% glycerol, 1 mM TCEP for Hsp90, Hop, Hsp40, Bag1, and p23, or 30 mM HEPES pH 7.5, 100 mM KCl, 10% glycerol, 1 mM TCEP for CHIP and Hsp70. Purified protein was concentrated, flash frozen, and stored at -80 degC.

### GRLBD expression and purification

For GR LBD, the ligand binding domain (LBD) (F602S) (520-777) was codon optimized and expressed in the pMAL-c3X derivative with an N-terminal cleavable 6x-His-MBP tag. GR LBD was expressed and purified as previously described^6,7^. GR DBD-LBD construct contains the GR DNA binding domain (DBD), hinge, and ligand binding domain (LBD) (418-777) with solubilizing mutation F602S. The construct was codon optimized and expressed in the pMAL-c3X derivative with an N-terminal cleavable 6xHis-MBP tag. GR DBD-LBD for cryo-EM sample preparation was expressed and purified as GR LBD. For the GR DBD-LBD used in ubiquitylation reactions in Fig. 2, purification was performed as previously described^14^.

### Assessing CHIP complexes in vitro

The GR chaperone cycle was reconstituted *in vitro* with separately purified components as previously described (Kirschke et al., 2014) in 30 mM Hepes pH 7.5, 50 mM KCl, 0.05% Tween-20, and 1 mM THP. To test for CHIP binding with GR alone, 5 uM MBP-GR LBD was incubated with 15 uM CHIP and 5 mM ATP/MgCl2. To test for CHIP binding with Hsp70-GR, 5 uM MBP-GR LBD was incubated with 2 uM Ydj1, 15 uM Hsp70, 15 uM CHIP, and 5 mM ATP/MgCl2. To test for CHIP binding with the GR-loading complex (Hsp90-GR-Hsp70-Hop), 5 uM MBP-GR LBD was incubated with 2 uM Ydj1, 15 uM Hsp70, 15 uM Hop, 15 uM D93N Hsp90, and 5 mM ATP/MgCl2. To test for CHIP binding with the GR-maturation complex (Hsp90-p23- GR), 5 uM MBP-GR LBD was incubated with 2 uM Ydj1, 5 uM Hsp70, 5 uM Hop, 15 uM Hsp90, 15 uM p23, and 5 mM ATP/MgCl2. The reaction was allowed to progress for 1 hour at room temperature. After 1 hr, 15 uM Bag-1, 20 uM CHIP, and 20 mM sodium molybdate (used to stabilize the closed conformation of Hsp90) were added, and the reaction was allowed to continue for another 30 minutes at room temperature.

Following incubation, reactions were added to amylose resin (New England Biolabs) in a 1:1 ratio and incubated at 4 degC with nutation for 1 hour. Resin was then washed 4 times with wash buffer and eluted with 50 mM maltose in elution buffer. To test for CHIP binding with GR alone, the wash buffer was 30 mM HEPES pH 7.5, 50 mM KCl, 0.05% Tween-20, 1 mM THP, and the elution buffer was 30 mM HEPES pH 7.5, 50 mM KCl, 50 mM maltose, 1 mM THP. To test for CHIP binding with Hsp70-GR as well as the GR-loading complex, the wash buffer was 30 mM HEPES pH 7.5, 50 mM KCl, 0.05% Tween-20, 400 uM ADP, 1 mM THP, and the elution buffer was 30 mM HEPES pH 7.5, 50 mM KCl, 400 uM ADP, 50 mM maltose, 1 mM THP. To test for CHIP binding with the GR-maturation complex, the wash buffer was 30 mM HEPES pH 7.5, 50 mM KCl, 0.05% Tween-20, 5 mM ATP, 20 mM sodium molybdate, 1 mM THP, and the elution buffer was 30 mM HEPES pH 7.5, 50 mM KCl, 20 mM sodium molybdate, 1 mM THP. Elutions were analyzed by SDS-PAGE.

### Hsp90-p23-GR-CHIP crosslinking analysis

For Fig. S2: CHIP was added to the *in vitro* reconstituted GR chaperone cycle using MBP-GR LBD, enriching the GR-maturation complex as described above. MBP-GR pulldown of Hsp90- p23-GR-CHIP was then injected over S200 3.2/300. Fractions B10, B9, and B8 from gel filtration containing Hsp90, p23, GR, and CHIP (as shown in Fig. 1d) were then treated with 0.05% glutaraldehyde and then visualized by SDS-PAGE and Western blot using anti-Hsp90 (Boston Biochemical), anti-CHIP (abcam), and anti-MBP (Rockland) antibodies.

### GR ubiquitylation

Ubiquitylation reactions were carried out in 30 mM Hepes pH 7.5, 50 mM KCl, 5 mM MgCl2, 1 mM THP. 100 nM MBP-GR DBD-LBD alone was incubated with varying concentrations of CHIP (50 nM, 25 nM, and 12.5 nM dimer) and 2 uM UbcH5c∼ub (Boston Biochemical) for 6 min. 100 nM MBP-GR DBD-LBD was pre-incubated with 15 uM Hsp70 and 2 uM Hsp40 in the presence of 5 mM ATP for 1 hr. The mixture was then incubated with varying concentrations of CHIP (50 nM, 25 nM, and 12.5 nM dimer) and 2 uM UbcH5c∼ub (Boston Biochemical) for 6 min. 100 nM Hsp90- p23-MBP-GR DBD-LBD (purified as above first by MBP-pulldown then size-exclusion chromatography via S200 3.2/300) was incubated with varying concentrations of CHIP (50 nM, 25 nM, and 12.5 nM dimer) and 2 uM UbcH5c∼ub (Boston Biochemical) for 6 min. Ubiquitylation reactions were quenched with 4X SDS buffer and 50 mM DTT and were visualized by Western blot using anti-MBP antibody (Rockland).

### Hsp90-p23-GR-CHIP sample preparation

To prepare the Hsp90-p23-GR-CHIP sample, CHIP was added to the *in vitro* reconstituted GR chaperone cycle using MBP-GR DBD-LBD, enriching the GR-maturation complex as described above. The elution from the MBP-GR affinity purification was treated with 0.05% glutaraldehyde for 20 minutes at room temperature. Crosslinked sample was purified by size exclusion using a Shodex KW-804 and fractions were analyzed by SDS-PAGE. Fractions containing the full complex was concentrated to ∼200 nM. 6.5 uL of sample was applied to PEG- 5k (polyethylene glycol 5000) carboxylate affinity grids (Wang et al., 2020) and plunge-frozen in liquid ethane using a Vitrobot Mark IV (FEI) with a wait time of 30 seconds, blotting time of 3 seconds, blotting force -10, at 8 degC, and with 100% humidity.

### Cryo-EM data acquisition

All data was acquired using SerialEM software. The dataset was collected on a Titan Krios G3 electron microscope (Thermo Fisher Scientific) operating at 300 kV using a K3 direct electron camera (Gatan). Images were recorded at a nominal magnification of 130,000x, corresponding to a physical pixel size of 0.652. The dataset was collected in non-super resolution mode. A nominal defocus range of 0.8 um - 2.0 um under focus was used for the dataset. The dose rate was 14.0 e-/pixel/second and the dose per frame was 0.82 e-/A2/frame, with 80 frames collected per movie. The total dose was 66 e-/A2.

### Cryo-EM data processing

The Hsp90-p23-GR-CHIP dataset consisted of ∼5500 dose-fractionated image stacks, which were motion corrected using UCSF MotionCor2^63^. Motion corrected images were used for contrast transfer function (CTF) estimation using Patch CTF Estimation in cryoSPARC (v.3.3.2) and then blob picking. 2D classification was performed on picks to remove high-resolution artifacts and iterated twice^64^. The particles were then exported from cryoSPARC using csparc2star^65^ to RELION (v. 4.0.0)^66^, where 3D classification took place using PDB 7KRJ as initial reference.

## Data availability

The cryo-EM maps generated in this study have been deposited in the Electron Microscopy Data Bank (EMDB) under the accession codes EMD-????? (Hsp90-p23-GR-CHIP) and EMD-????? (apo Hsp90-p23-GR).

## References

1. Nathan, D. F., & Lindquist, S. Mutational analysis of Hsp90 function: interactions with a steroid receptor and a protein kinase. Mol Cell Biol, 15, 3917–3925, doi:10.1128/mcb.15.7.3917 (1995).

2. Taipale, M., Krykbaeva, I., Koeva, M., Kayatekin, C., Westover, K. D., Karras, G. I., & Lindquist, S. Quantitative analysis of HSP90-client interactions reveals principles of substrate recognition. Cell, 150, 987–1001, doi:10.1016/j.cell.2012.06.047 (2012).

3. Cyr, D., Hohfeld, J., & Patterson, C. Protein quality control: U-box-containing E3 ubiquitin ligases join the fold. Trends Biochem Sci, 27, 368–375, doi:10.1016/s0968-0004(02)02125-4 (2002).

4. McClellan, A. J., Tam, S., Kaganovich, D., & Frydman, J. Protein quality control: chaperones culling corrupt conformations. Nat Cell Biol, 7, 736–741, doi:10.1038/ncb0805-736 (2005).

5. Chen, B., Retzlaff, M., Roos, T., & Frydman, J. Cellular strategies of protein quality control. Cold Spring Harb Perspect Biol, 3, a004374, doi:10.1101/cshperspect.a004374 (2011).

6. Kirschke, E., Goswami, D., Southworth D., Griffin, P., & Agard, D. A. Glucocorticoid receptor function regulated by coordinated action of the Hsp90 and Hsp70 chaperone cycles. Cell, 157, 1685–1697, doi:10.1016/j.cell.2014.04.038 (2014).

7. Noddings, C. M., Wang, R. Y.-R., Johnson, J. L., & Agard, D. A. Structure of Hsp90-p23- GR reveals the Hsp90 client-remodeling mechanism. Nature, 601, 465–469, doi:10.1038/s41586-021-04236-1 (2022).

8. Hartl, F. U., Bracher, A., & Hayer-Hartl, M. Molecular chaperones in protein folding and proteostasis. Nature, 475, 324–332. doi:10.1038/nature10317 (2011).

9. Gottesman, S., Wickner, S., Maurizi, M. R. Protein quality control: triage by chaperones and proteases. Genes Dev, 11, 815–823, doi:10.1101/gad.11.7.815 (1997).

10. Wickner, S., Maurizi, M. R., Gottesman, S. Posttranslational quality control: folding, refolding, and degrading proteins. Science, 286, 1888–1893, doi:10.1126/science.286.5446.1888 (1999).

11. Shao, S., Rodrigo-Brenni, M. C., Kivlen, M. H., Hegde, R. S. Mechanistic basis of a molecular triage reaction. Science, 355, 298–302, doi:10.1126/science.aah6130 (2017).

12. Verba, K. A., Wang, R. Y. R, Arakawa, A, Liu, Y., Shirouzu, M., Yokoyama, S., & Agard, D. A. Atomic structure of Hsp90-Cdc37-Cdk4 reveals that Hsp90 traps and stabilizes an unfolded kinase. Science, 352, 1542–1547, doi:10.1126/science.aaf5023. (2016).

13. Wang, R. Y. R., Noddings, C. M., Kirschke, E., Myasnikov, A. G., Johnson, J. L., & Agard, D. A. Structure of Hsp90-Hsp70-Hop-GR reveals the Hsp90 client-loading mechanism. Nature, 601, 460–464, doi:10.1038/s41586-021-04252-1 (2022).

14. Noddings, C. M., Johnson, J. L., & Agard, D. A. Cryo-EM reveals how Hsp90 and FKBP immunophilins co-regulate the glucocorticoid receptor. Nat Struct Mol Biol, 30, 1867–1877, doi:10.1038/s41594-023-01128-y (2023).

15. Oberoi, J., Guiu, X. A., Outwin, E. A., Schellenberger, P., Roumeliotis, T. I., Choudhary, J. S., & Pearl, L. H. HSP90-CDC37-PP5 forms a structural platform for kinase dephosphorylation. Nat Comm, 13, 7343, doi:10.1038/s41467-022-35143-2 (2022).

16. Jaime-Garza, M., Nowotny, C. A., Coutandin, D., Wang, F., Tabios, M., & Agard. D. A. Hsp90 provides a platform for kinase dephosphorylation by PP5. Nat Commun, 14, 2197, doi:10.1038/s41467-023-37659-7 (2023).

17. Finci, L. I., Chakrabarti, M., Gulten, G., Finney, J., Grose, C., Fox, T., Yang, R., Nissley, D. V., McCormick, F., Esposito, D., Balius, T. E., & Simanshu, D. K. Structural dynamics of RAF1-HSP90-CDC37 and HSP90 complexes reveal asymmetric client interactions and key structural elements. Commun Biol 7, 260, doi:10.1038/s42003-024-05959-3 (2024).

18. Neckers, L. Hsp90 inhibitors as novel cancer chemotherapeutic agents. Trends Mol Med., 8, S55–61, doi:10.1016/s1471-4914(02)02316-x (2002).

19. Whitesell, L., Mimnaugh, E. G., De Costa, B., Myers, C. E., & Neckers, L. M. Inhibition of heat shock protein HSP90-pp60v-src heteroptotein complex formation by benzoquinone ansamycins: essential role for stress proteins in oncogenic transformation. Proc Natl Acad Sci USA, 91, 8324–8328, doi: 10.1073/pnas.91.18.8324 (1994).

20. Miller, P., DiOrio, C., Moyer, M., Schnur, R. C., Bruskin, A., Cullen, W., & Moyer, J. D. Depletion of the erbB-2 gene product p185 by benzoquinoid ansamycins. Cancer Res, 54, 2724–30 (1994).

21. Blagoskonny, M. V., Toretsky, J., Bohen, S., & Neckers, L. Mutant conformation of p53 translated in vitro or in vivo requires functional HSP90. Proc Natl Acad Sci USA, 93, 8379–8383, doi:10.1073/pnas.93.16.8379 (1996).

22. Mimnaugh, E. G., Chavany, C., & Neckers, L. Polyubiquitination and proteasomal degradation of the p185c-erbB-2 receptor protein-tyrosine kinase induced by geldamycin. J Biol Chem, 271, 22796–801, doi:10.1074/jbc.271.37.22796 (1996).

23. Schulte, T. W., Blagosklonny, M. V., Romanova, L., Mushinski, J. F., Monia, B. P., Johnston, J. F., Nguyen, P., Trepel, J., & Neckers, L. Destabilization of Raf-1 by geldanamycin leads to disruption of the Raf-1-MEK-mitogen-activated protein kinase signaling pathway. Mol Cell Biol, 16, 5839–5845, doi:10.1128/MCB.16.10.5839 (1996).

24. Whitesell, L., & Cook, P. Stable and specific binding of heat shock protein 90 by geldanamycin disrupts glucocorticoid receptor function in intact cells. Mol Endrocrinol, 10, 705–712, doi:10.1210/mend.10.6.8776730 (1996).

25. Segnitz, B., & Gehring, U. The function of steroid hormone receptors is inhibited by the hsp90-specific compound geldanamycin. J Biol Chem, 272, 18694–18701, doi: 10.1074/jbc.272.30.18694 (1997).

26. Connell, P., Ballinger, C. A., Jiang, J., Wu, Y., Thompson, L. J., Hohfeld, J., & Patterson, C. The co-chaperone CHIP regulates protein triage decisions mediated by heat-shock proteins. Nat Cell Biol, 3, 93–96, doi:10.1038/35050618 (2001).

27. Xu, W., Marcu, M., Yuan, X., Mimnaugh, E., Patterson, C., & Neckers, L. Chaperone-dependent E3 ubiquitin ligase CHIP mediates a degradative pathway for c-ErbB2/Neu. Proc Natl Acad Sci USA, 99, 12847–12852, doi:10.1073/pnas.202365899 (2002).

28. Dickey, C. A., Kamal, A., Lundgren, K., Klosak, N., Bailey, R. M., Dunmore, J., Ash, P., Shoraka, S., Zlatkovic, J., Eckman, C. B., Patterson, C., Dickson, D. W., Nahman Jr., N. S., Hutton, M., Burrows, F., & Petrucelli, L. The high-affinity HSP90-CHIP complex recognizes and selectively degrades phosphorylated tau client proteins. J Clin Invest, 117, 648–658, doi:10.1172/JCI29715 (2007).

29. Ehrlich, E. S., Wang, T., Luo, K., Xiao, Z., Niewiadomska, A. M., Martinez, T., Xu, W., Neckers, L., Yu, X.-F. Regulation of Hsp90 client proteins by a Cullin5-RING E3 ubiquitin ligase. Proc Natl Acad Sci USA, 106, 20330–20335, doi:10.1073/pnas.0810571106 (2009).

30. Samant, R. S., Clarke, P. A., Workman, P. E3 ubiquitin ligase Cullin-5 modulates multiple molecular and cellular responses to heat shock protein 90 inhibition in human cancer cells. Proc Natl Acad Sci USA, 111, 6834–6839, doi:10.1073/pnas.1322412111 (2014).

31. Taipaile, M., Tucker, G., Peng, J., Krybaeva, I., Lin, Z.-Y., Larsen, B., Choi, Y., Berger, B., Gingras, A.-C., & Lindquist, S. Cell 158, 434–448, doi:10.1016/j.cell.2014.05.039 (2014).

32. Li, Z., Zhou, L., Prodromou, C., Savic, V., & Pearl, L. H. (2017). HECTD3 mediates an HSP90-dependent degradation pathway for protein kinase clients. Cell Rep, 19, 2515–2528, doi:10.1016/j.celrep.2017.05.078 (2017).

33. Hershko A., & Ciechanover, A. The ubiquitin system. Annu Rev Biochem, 67, 425–479, doi:10.1146/annurev.biochem.67.1.425 (1998).

34. Finley, D. Recognition and processing of ubiquitin-protein conjugates by the proteasome. Annu Rev Biochem, 78, 477–513, doi:10.1146/annurev.biochem.78.081507.101607 (2009).

35. Tsuboyama, K., Tadakuma, H., & Tomari, Y. Conformational activation of Argonaute by distinct yet coordinated actions of the Hsp70 and Hsp90 chaperone systems. Mol Cell, 70, 722–729, doi:10.1016/j.molcel.2018.04.010 (2018).

36. Boysen, M., Kityk, R., & Mayer, M. P. Hsp70- and Hsp90-mediated regulation of the conformation of p53 DNA binding domain and p53 cancer variants. Mol Cell, 74, 831–843, doi:10.1016/j.molcel.2019.03.032 (2019).

37. Dahiya, V., Agam, G., Lawatscheck, J., Rutz, D. A., Lamb, D. C., & Buchner, J. Coordinated conformational processing of the tumor suppressor protein p53 by the Hsp70 and Hsp90 chaperone machineries. Mol Cell, 74, 816–830, doi:10.1016/j.molcel.2019.03.026 (2019).

38. Ballinger, C. A., Connell, P., Wu, Y., Hu, Z., Thompson, L. J., Yin, L. Y., & Patterson, C. Identification of CHIP, a novel tetratricopeptide repeat-containing protein that interacts with heat shock proteins and negatively regulates chaperone functions. Mol Cell Biol, 19, 4535–4545, doi:10.1128/MCB.19.6.4535 (1999).

39. Hatakeyama, S., Yada, M., Matsumoto, M., Ishida, N., & Nakayama, K. I. U box proteins as a new family of ubiquitin-protein ligases. J Biol Chem, 276, 33111–33120, doi:10.1074/jbc.M102755200 (2001)

40. Connell, P., Ballinger, C. A., Jiang, J., Wu, Y., Thompson, L. J., Hohfeld, J., & Patterson, C. The co-chaperone CHIP regulates protein triage decisions mediated by heat-shock proteins. Nat Cell Biol 3, 93–96, doi:10.1038/35050618 (2001).

41. Demand, J., Alberti, S., Patterson, C., & Hohfeld, J. Cooperation of a ubiquitin domain protein and an E3 ubiquitin ligase during chaperone/proteasome coupling. Curr Biol, 11, 1569–1577, doi:10.1016/s0960-9822(01)00487-0 (2001).

42. Stankiewicz, M., Nikolay, R., Rybin, V., & Mayer, M. P. CHIP participates in protein triage decisions by preferentially ubiquitinating Hsp70-bound substrates. FEBS J, 277, 3353–3367, doi:10.1111/j.1742-4658.2010.07737.x (2010).

43. Esser, C., Scheffner, M., & Hohfeld, J. The chaperone-associated ubiquitin ligase CHIP is able to target p53 for proteasomal degradation. J Biol Chem, 280, 27443–27448, doi:10.1074/jbc.M501574200 (2005).

44. Ren, H. Y., Patterson, C., Cyr, D. M., & Rosser, M. F. N. Reconstitution of CHIP E3 ubiquitin ligase activity. Methods Mol Biol, 787, 93–103, doi:10.1007/978-1-61779-295-3_8 (2011).

45. Younger, J. M., Ren, H.-Y., Chen, L., Fan, C.-Y., Fields, A., Patterson, C., & Cyr, D. M. A foldable CFTRΔF508 biogenic intermediate accumulates upon inhibition of the Hsc70- CHIP E3 ubiquitin ligase. J Cell Biol, 167, 1075–1085, doi:10.1083/jcb.200410065 (2004).

46. Murata, S., Minami, Y., Minami, M., Chiba, T., & Tanaka, K. (2001). CHIP is a chaperone-dependent E3 ligase that ubiquitylates unfolded protein. EMBO Rep, 2, 1133–1138, doi: 10.1093/embo-reports/kve246 (2001).

47. Quintana-Gallardo, L., Martin-Benito, J., Marcilla, M., Espadas, G., Sabido, E., & Valpuesta, J. M. The cochaperone CHIP marks Hsp70- and Hsp90-bound substrates for degradation through a very flexible mechanism. Sci Rep 9, 5102, doi:10.1038/s41598-019-41060-0 (2019).

48. Kirschke, E., Roe-Zurz, Z., Noddings, C., & Agard, D. The interplay between Bag-1, Hsp70, and Hsp90 reveals that inhibiting Hsp70 rebinding is essential for glucocorticoid receptor activity. bioRxiv, doi:10.1101/2020.05.03.075523 (2020).

49. Johnson, J. L. & Toft, D. O. Binding of p23 and hsp90 during assembly with the progesterone receptor. Mol Endocrinol 9, 670–678, doi:10.1210/mend.9.6.8592513 (1995).

50. Ali, M. M. U., Roe, S. M., Vaughan, C. K., Meyer, P., Panaretou, B., Piper, P. W., Prodromou, C., & Pearl, L. H. Crystal structure of an Hsp90-nucleotide-p23/Sba1 closed chaperone complex. Nature 440, 1013–1017, doi:10.1038/nature04716. (2006).

51. Pickart, C. M., & Eddins, M. J. Ubiquitin: structures, functions, mechanisms. Biochim Biophys Acta, 1695, 55–72, doi:10.1016/j.bbamcr.2004.09.019 (2004).

52. Pickart, C. M. Mechanisms underlying ubiquitination. Annu Rev Biochem 70, 503–533, doi:10.1146/annurev.biochem.70.1.503 (2001).

53. Wang, F., Yu, Z., Betegon, M., Campbell, M. G., Aksel, T., Zhao, J., Li, S., Douglas, S. M., Cheng. Y., & Agard, D. A. Amino and PEG-amino graphene oxide grids enrich and protect samples for high-resolution single particle cryo-electron microscopy. J Struct Biol 209, 107437, doi:10.1016/j.jsb.2019.107437 (2020).

54. Zhang, M., Windheim, M., Roe, S. M., Peggie, M., Cohen, P., Prodromou, C., & Pearl, L. H. Chaperoned ubiquitylation—crystal structures of the CHIP U box E3 ubiquitin ligase and a CHIP-Ubc13-Uev1a complex. Mol Cell 20, 525–538, doi:10.1016/j.molcel.2005.09.023 (2005).

55. Balaji, V., Muller, L., Lorenz, R., Kevei, E., Zhang, W. H., Santiago, U., Gebauer, J., Llamas, E., Vilchez, D., Camacho, C. J., Pokrzywa, W., & Hoppe, T. A dimer-monomer switch controls CHIP-dependent substrate ubiquitylation and processing. Mol Cell 82, 3239–3254, doi:10.1016/j.molcel.2022.08.003, (2022).

56. Bledsoe, R. K., Montana, V. G., Stanley, T. B., Delves, C. J., Apolito, C. J., McKee, D. D., Consler, T. G., Parks, D. J., Stewart, E. L., Willson, T. M., Lambert, M. H., Moore, J. T., Pearce, K. H., & Xu, H. E. Crystal structure of the glucocorticoid receptor ligand binding domain reveals a novel mode of receptor dimerization and coactivator recognition. Cell 110, 93–105, doi:10.1016/s0092-8674(02)00817-6 (2002).

57. Bourget, W., Ruff, M. Chambon, P. Gronemeyer, H., & Moras, D. Crystal structure of the ligand-binding domain of the human nuclear receptor RXR-alpha. Nature 375, 377–382, doi:10.1038/375377a0 (1995).

58. Nolte, R. T., Wisely, G. B., Westin, S., Cobb, J. E., Lambert, M. H., Kurokawa, R., Rosenfeld, M. G., Willson, T. M., Glass, C. K., & Milburn, M. V. Ligand binding and co- activator assembly of the peroxisome proliferator-activated receptor-. Nature 395, 137–143, doi:10.1038/25931 (1998).

59. Watkins, R. E., Wisely, G. B., Moore, L. B., Collins, J. L., Lambert, M. H., Williams, S. P., Willson, T. M., Kliewer, S. A., & Redinbo, M. R. The human nuclear xenobiotic receptor PXR: structural determinants of directed promiscuity. Science 292, 2329–2333, doi:10.1126/science.1060762 (2001).

60. Noguchi, M., Nomura, A., Murase, K., Doi, S., Yamaguchi, K., Hirata, K., Shiozaki, M., Hirashima, S., Kotoku, M., Yamaguchi, T., Katsuda, Y., Steensma, R., Li, X., Tao, H., Tse, B., Fenn, M., Babine, R., Bradley, E., Crowe, P., Thacher, S., Adachi, T., & Kamada, M. Ternary complex of human RORγ ligand-binding domain, inverse agonist and SMR peptide shows a unique mechanism of corepressor recruitment. Genes Cells 22, 535–551, doi:10.1111/gtc.12494 (2017).

61. Li, X., Anderson, M., Collin, D., Muegge, I., Wan, J., Brennan, D., Kugler, S., Terenzio, D., Kennedy, C., Lin, S., Labadia, M. E., Cook, B., Hughes, R., & Farrow, N. A. Structural studies unravel the active conformation of apo RORγt nuclear receptor and a common inverse agonism of two diverse classes of RORγt inhibitors. J Biol Chem 292, 11618–11630, doi:10.1074/jbc.M117.789024 (2017).

62. Suren, T., Rutz, D., Mößmer, P., Merkle, U., Buchner, J., & Rief, M. Single-molecule force spectroscopy reveals folding steps associated with hormone binding and activation of the glucocorticoid receptor. Proc Natl Acad Sci USA 115, 11688–11693, doi:10.1073/pnas.1807618115 (2018).

63. Zheng, S. Q., Palovcak, E., Armache, J.-P., Verba, K. A., Cheng, Y., & Agard, D. A. MotionCor2: anisotropic correction of beam-induced motion for improved cryo-electron microscopy. Nat Methods 14, 331–332, doi:10.1038/nmeth.4193 (2017).

64. Punjani, A., Rubinstein, J. L., Fleet, D. J., & Brubaker, M. A. cryoSPARC: algorithms for rapid unsupervised cryo-EM structure determination. Nat Methods 14, 290–296, doi:10.1038/nmeth.4169 (2017).

65. Asarnow, D., Palovcak, E., & Cheng, Y. UCSF pyem v0.5. Zenodo, 10.5281/zenodo3576630 (2019)

66. Kimanius, D., Dong, L., Sharov, G., Nakane, T., & Scheres, S. H. W. New tools for automated cryo-EM single-particle analysis in RELION-4.0. Biochem J 478, 4169–4185, doi:10.1042/BCJ20210708 (2021).

